# Quantitative cross-species translators of cardiac myocyte electrophysiology: model training, experimental validation, and applications

**DOI:** 10.1101/2020.12.17.423297

**Authors:** Stefano Morotti, Caroline Liu, Bence Hegyi, Haibo Ni, Alex Fogli Iseppe, Lianguo Wang, Crystal M. Ripplinger, Donald M. Bers, Andrew G. Edwards, Eleonora Grandi

## Abstract

Animal experimentation is key in the evaluation of cardiac efficacy and safety of novel therapeutic compounds. However, inter-species differences in the mechanisms regulating excitation-contraction coupling can limit the translation of experimental findings from animal models to human physiology, and undermine the assessment of drugs’ efficacy and safety. Here, we built a suite of translators for quantitatively mapping electrophysiological responses in ventricular myocytes across species. We trained these statistical operators using a broad dataset obtained by simulating populations of our biophysically detailed computational models of action potential and Ca^2+^ transient in mouse, rabbit, and human. We then tested our translators against experimental data describing the response to stimuli, such as ion channel block, change in beating rate, and β-adrenergic challenge. We demonstrate that this approach is well suited to predicting the effects of perturbations across different species or experimental conditions, and suggest its integration into mechanistic studies and drug development pipelines.

## Introduction

Cardiovascular disease is the leading cause of morbidity and mortality worldwide.^1,2^ The combined efforts of basic, translational, and clinical research have greatly augmented our understanding of disease mechanisms over recent decades, but many challenges still exist. Among these are the problems associated with interpreting in the context of human disease preclinical experimentation conducted in animals, and in various *in vitro* and animal models of disease.^3,4^ Physiological species-differences present a major intrinsic limitation to translating those experimental findings from animals to human. Given the wide adoption of animal experimentation in the processes involved in the development of therapeutics, implications of inter-species differences are particularly important in the pharmaceutical field, where there are growing concerns about safety and efficacy of drugs tested on animals.^5,6^

The species most commonly used for preclinical assessment of cardiac electrophysiologic outcomes are small mammals,^7,8^ including mice^9^ and rabbits.^10^ Despite genetic similarities, differences in cardiac function among mammals are evident at both organ and cellular level. For example, body and heart weights, as well as stroke volume, vary across approximately three orders of magnitude, and resting heart rate is about ten-fold higher in mouse vs. human (~600 vs. 60 bpm).^9^ To accommodate for these quite different working regimes, evolution has led to several differences in the ionic mechanisms controlling excitation-contraction coupling (ECC).^7,11^ Specifically, varying expression and regulation of ion channels and transporters, notably K^+^ channels,^12–15^ is mechanistically associated with species-specific action potential (AP) properties. It has been shown, for example, that the presence of a more prominent “spike & dome” morphology is due to a large transient outward current I_to_ in species like rabbit and human, while I_to_ is virtually absent in the guinea pig, which lacks the AP notch.^16^ A much larger I_to_ and expression of additional K^+^ channels are responsible for the typical triangular AP shape and shorter AP duration (APD) in mouse and rat (vs. non-rodents) ventricular myocytes.^17^ A notable implication of these dissimilarities is that the same perturbation (e.g., drug administration) can lead to markedly different changes in the AP and intracellular Ca^2+^ transient (CaT) properties in different species. This can occur even in species with comparable AP profiles, for example, due to selective block of the slow delayed rectifying K^+^ current (I_Ks_), which prolongs AP in guinea pig but does not significantly alter rabbit or human APD.^18–20^ Moreover, these differences, even when relatively subtle, can markedly alter propensity for arrhythmogenic voltage and Ca^2+^ instabilities.^13^ Recent research has also highlighted inter-species differences in the APD changes leading to optimal inotropic response to β-adrenergic receptor (β-AR) stimulation,^21,22^ a well-known mediator of cardiac stress responses and major arrhythmia trigger.^23^ Thus, significant differences in ECC regulation in human vs. animal models have important implications for the study of arrhythmogenic mechanisms and, consequently, for the development of pharmacological anti-arrhythmic treatments in patients.

With the recent development and widespread use of experimental technologies based on induced pluripotent stem cells (iPSCs), human iPSC-derived cardiomyocytes (hiPSC-CMs) have become a common alternative to animal models in cardiovascular and pharmaceutical research.^24^ Since hiPSC-CMs retain the genetic information of the donors they are derived from, these cells are an ideal tool for the investigation of patient-specific physiology and response to drugs.^25,26^ However, the high degree of variability in electrophysiological properties among different hiPSC-CM lines, and within the same culture,^27^ and the immature phenotype characterized by altered expression of ion channels, spontaneous beating activity, and impaired contractility, can limit translation of responses in hiPSC-CMs to adult cardiac myocytes.^28^ To address this limitation, Gong & Sobie recently proposed an approach that combines simulations and statistical analysis to create quantitative “translators” that map electrophysiological responses across different cell types.^29^ Their work theoretically demonstrated the feasibility of regression-based operators that take as input experimental data recorded in hiPSC-CMs, and directly produce as output the predicted effect in adult myocytes,^29^ and suggested translation across species is also achievable using existing models. Here, we coupled the Gong & Sobie regression-based approach with our established lineage of ventricular myocyte AP and CaT models in mouse,^30,31^ rabbit,^32,33^ and human,^34,35^ and constructed a suite of predictors of human electrophysiological response from mouse and rabbit data. We validated our translators against a broad set of experimental data, and demonstrated their suitability to predict human response to pharmacological perturbation from experiments in animal models, suggesting that their systematic integration into the drug development pipeline could facilitate the assessment of efficacy and safety of novel compounds.

## Results

### Sensitivity analysis of multi-species models reveals species differences in ECC properties

We updated our established models of ventricular myocytes in mouse,^30,31^ rabbit,^32,33^ and human,^34,35^ and created a coherent multi-species computational framework for ECC simulations. This suite of models integrates detailed description of membrane electrophysiology, intracellular Ca^2+^ and Na^+^ handling, and Ca^2+^/calmodulin-dependent protein kinase II (CaMKII) and β-AR signaling cascades. It also includes detailed characterization of species-specific electrophysiological properties recapitulating well-known species differences in AP profile and Ca^2+^ handling (**Fig. 1A** & **Fig. S1**). Following an established approach (**Fig. 1B**),^36^ we created populations of 1,000 mouse, rabbit, and human myocyte models that replicate the natural cell-to-cell variability seen in experiments (**Fig. 1C**) by randomly perturbing baseline model parameters (defined in **Table S1**). We then assessed steady-state AP and CaT features (defined in **Table 1**) for each model variant in the three populations, and performed multivariable linear regression to quantify the sensitivity of such features to changes in perturbed parameters.^36^ This systematic analysis shows that APD and CaT are differently regulated in mouse, rabbit, and human ventricular myocytes (**Fig. 1D** & **Fig. S2**). AP repolarization, similarly in rabbit and human, is mostly controlled by I_to_, I_Ks_, and the rapid delayed rectifying K^+^ current (I_Kr_). The shorter and more triangular murine AP is more sensitive to changes in the inwardly rectifying K^+^ current (I_K1_), and strongly affected by changes in the mouse-specific ultra-rapidly activating and slowly inactivating (I_K,slow_), and non-inactivating steady-state (I_ss_) K^+^ currents. These results demonstrate that a similar perturbation (e.g., selective ion channel block) can cause APD and CaT changes that are quantitatively different among different species.

**Table 1.**
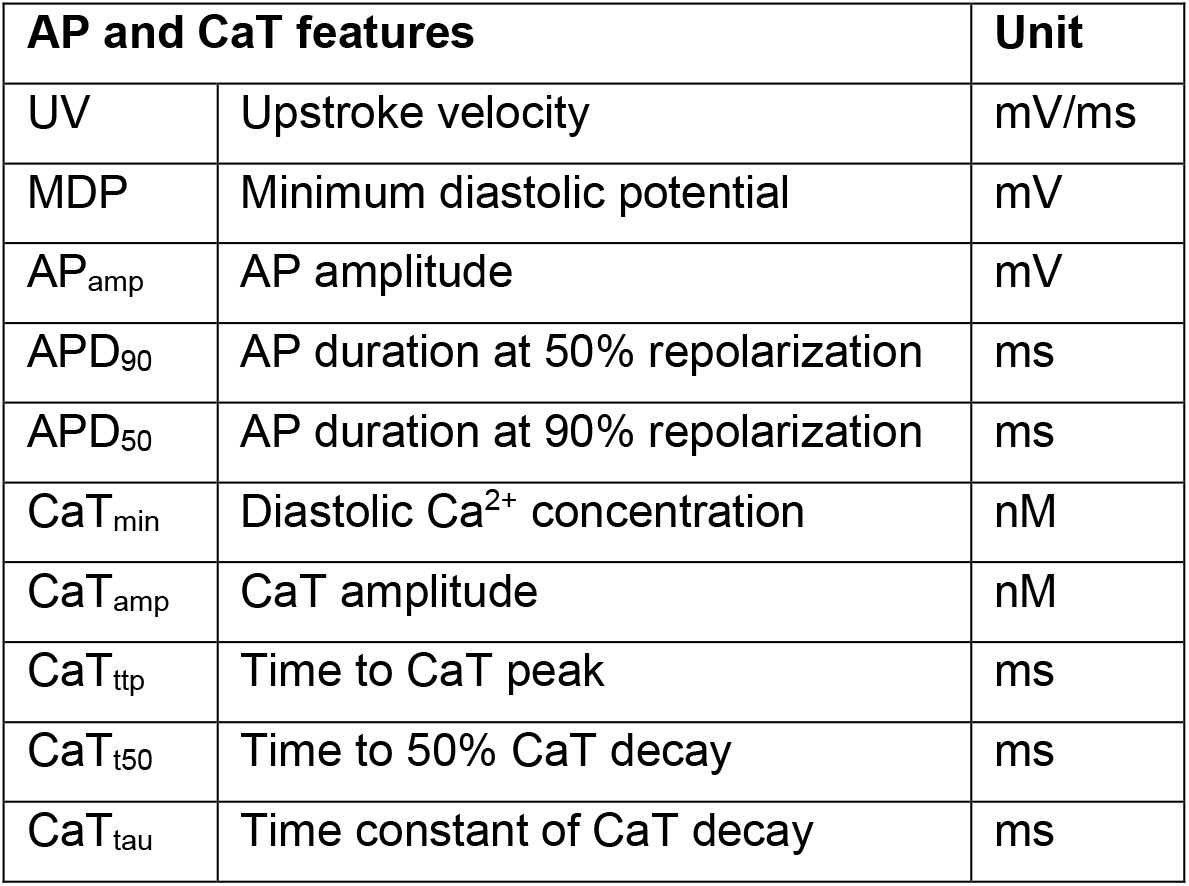
Definition of action potential and Ca^2+^ transient features.

**Figure 1.**
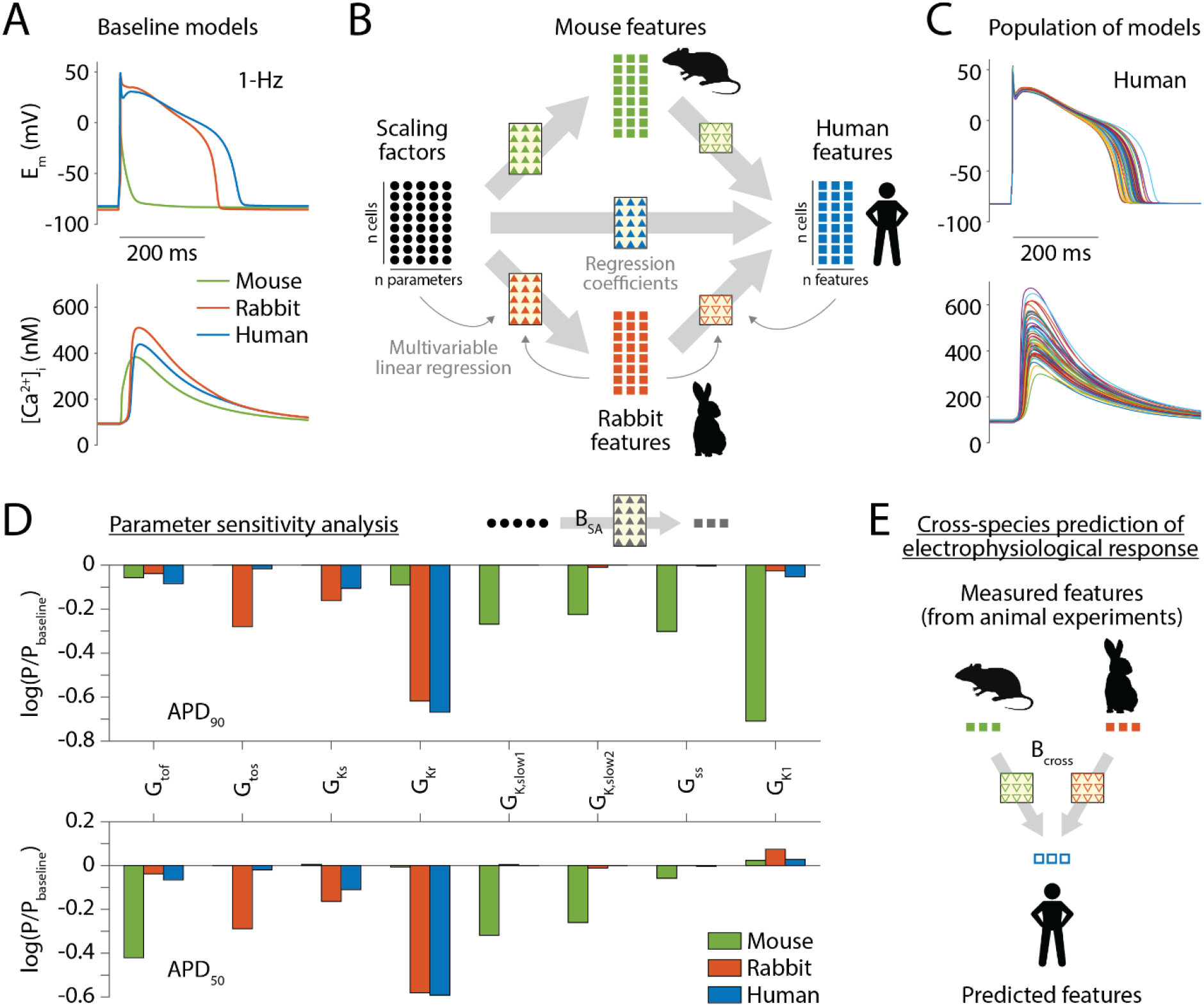
Overview of the approach. **A**) Simulated AP and CaT traces elicited by stimulating the baseline mouse, rabbit, and human models at 1 Hz in control condition. **B**) The same matrix of randomly generated scaling factors (represented with circles) is used to perturb the values of selected parameters in the baseline mouse, rabbit, and human models, thus creating three populations of models. Panel **C** highlights the variability among 100 representative AP and CaT traces within a population of human models (1-Hz pacing, control). AP and CaT features (squares) are estimated at steady-state in each model variant in the three populations, and their values collected in three matrices. Multivariable regression analysis between the matrix of scaling factors and the matrix of features is performed to assess the sensitivity of AP and CaT features to changes in model parameters in the three species. The result of this process is the regression matrix B_SA_, which coefficients (triangles) can be used to estimate the values of AP and CaT features given a known set of parameter scaling factors. Panel **D** displays the regression coefficients quantifying the sensitivity of APD_90_ and APD_50_ to changes in the maximal conductance of repolarizing K^+^ channels in the three species (1-Hz pacing, control). Multivariable regression analysis between the features matrices in two different species is performed to obtain the cross-cell type translation matrix B_cross_ that can be used to predict AP and CaT features in the output species, given the values observed in the input species. Panel **E** shows the proposed application of cross-cell type translators (mouse-to-human and rabbit-to-human) to predict human response from experimental findings in animal models.

### Multivariable linear regression is used to build translators of ECC properties across species or experimental conditions

Using the approach proposed by Gong & Sobie,^29^ we built populations of models by perturbing only the parameters that are common to the mouse, rabbit, and human models, and applied multivariable linear regression between sets of species-specific AP and CaT features to generate a suite of statistical translators (a set of regression coefficients) for mapping mouse and rabbit data onto human physiologic responses (**Fig. 1B**). Upon validation, these mouse-to-human and rabbit-to-human translators could be directly applied to predict human features from data obtained from experiments in animal models (**Fig. 1E**).

Examples of cross-species translators are shown in **Fig. 2A-B**. Each predictor was built using four simulated features for both input and output species, namely APD at 50 and 90% of repolarization (APD_50_ and APD_90_), time to 50%, and time constant of CaT decay (CaT_t50_ and CaT_tau_), from populations of *in silico* cells. A given output feature (e.g., APD_90_ in human) is calculated by applying a function in which each one of four input features in mouse or rabbit is multiplied by the corresponding regression coefficient (i.e., the four values shown in one row in the matrix).^29^ We then tested these translators using simulated data from independent populations of models, i.e., comparing the actual (simulated) to the predicted (translated) features (**Fig. 2A-B**, scatter plots). Results of validation show that human CaT duration measurements could be well reproduced translating both mouse and rabbit data, while prediction of human APD data from mouse is less accurate than starting from rabbit data. Nevertheless, predictions of human APD values from mouse data are remarkably close to the actual data if considering the distance “traveled” between input and output (**Fig. S3A**).

**Figure 2.**
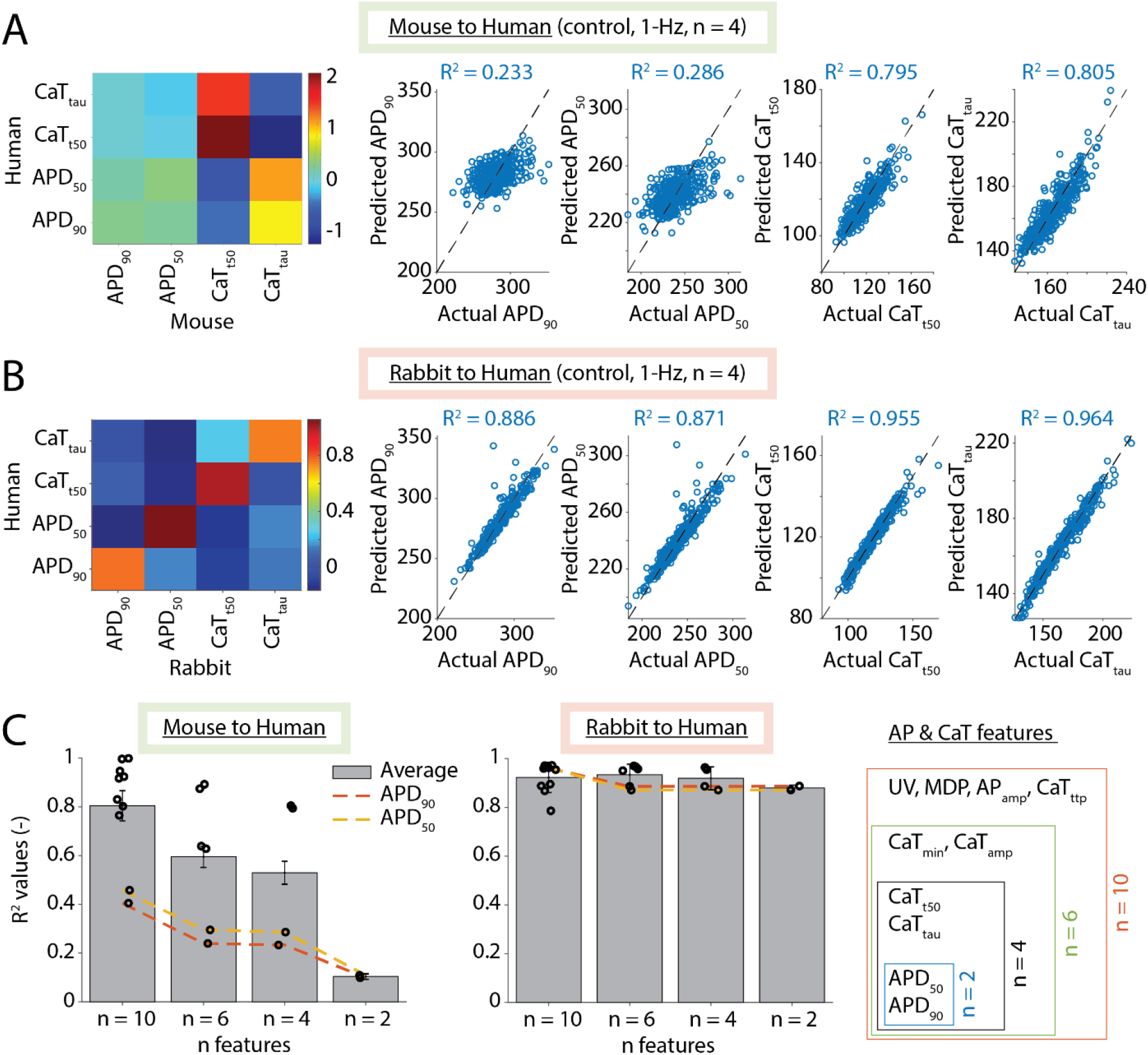
Development and validation of cross-species translators of electrophysiological response. Mouse-to-human (**A**) and rabbit-to-human (**B**) translation matrices B_cross_ are built using mouse (or rabbit) and human features (APD_90_, APD_50_, CaT_t50_, and CaT_tau_) assessed in population of models paced at 1 Hz (control condition). Scatter plots on the right in panels **A** and **B** show the result of validation performed with an independent set of simulated data, obtained in different populations counting 400 models. For each feature, we plot predicted human features (obtained by applying B_cross_ to the data produced simulating the input species, ordinate) against the actual values from human simulations (abscissa), and indicate the coefficient of determination R^2^. **C**) Bar graphs show the mean R^2^ values obtained averaging the R^2^ values estimated for each feature by predictors built using a varying number *n* of both input and output features (as illustrated by the schematic on the right). Circles correspond to the R^2^ values for each feature, and error bars indicate standard deviation of the mean.

We investigated the translators’ performance when varying the number and composition of input and output features (**Fig. 2C**). We compared the overall performance (estimated as average R^2^ value) and found that prediction from rabbit is always more accurate than prediction from mouse. Starting from a full set of ten AP and CaT features (**Table 1**), we progressively eliminated features to mimic more realistic conditions in which experiments offer a less rich dataset. As expected, reducing the number of features worsens the accuracy of the translation. This effect is especially evident for mouse-to-human translation, where impaired APD prediction negatively impacts the overall performance. Moreover, prediction of human APD values (dashed lines in **Fig. 2C**) from mouse data improves when it relies on other input features (other than APD_50_ and APD_90_). Performance of rabbit-to-human translation is minimally affected by the reduction of features, and APD predictions of the “minimal” translator (*n* = 2) are still comparable with those obtained using the whole dataset.

### Cross-species translators reliably predict species-specific simulated effects of ion current blockade

Although the accuracy of human prediction from rabbit data is generally better than the one from mouse (**Fig. 2C**), human features predicted from both mouse and rabbit are close to the actual simulated data if considering the distance between input and output values. This can be appreciated by considering the results of cross-species translation of features obtained from simulating our baseline models in control condition (**Fig. S3A**). To extend the characterization of translators’ performance, we tested their ability to translate the effects of simulated drug application. AP and CaT values obtained through simulation of selective ion channel block in our baseline mouse and rabbit models were translated into human using the previously described predictors (i.e., those developed using control data).

Response to drug is qualitatively similar in rabbit and human (**Fig. 3A**), and predictions from rabbit match human simulated data quite well (**Fig. 3B-G**). Generally, minimal rabbit-to-human translators produce reliable predictions, and increasing *n* does not lead to any appreciable improvement. Mouse-to-human translators lead to less and more variably accurate predictions, where best results were obtained for translating the effects of blocking the fast component of I_to_ (I_tof_) and the L-type Ca^2+^ current (I_CaL_). In these cases, increasing *n* improved the prediction, especially for CaT measurements. Changes in APD_90_ and CaT_tau_ were also well reproduced when predicting the effects induced by block of I_K1_ and late Na^+^ current (I_NaL_), while the worst predictions were obtained for I_Kr_ block. The latter result is not surprising, because I_Kr_ plays a major role in shaping AP repolarization in larger mammals, but minimally affects AP (or CaT) in mice (**Fig. S2**). Our mouse-to-human translators also failed to accurately predict response to Na^+^/Ca^2+^ exchanger (NCX) block because of the development of arrhythmogenic spontaneous Ca^2+^ release events from the sarcoplasmic reticulum (SR) in mouse simulations (**Fig. S4**). For both mouse-to-human and rabbit-to-human translations, predicted APD_50_ and CaT_t50_ values show a trend similar to those of APD_90_ and CaT_tau_, respectively (**Fig. S3B-F**). Taken together, our data show that prediction of human responses from rabbit data is generally accurate even when only a few measurements are available. On the other hand, prediction from mouse data is more challenging because of intrinsic differences in the regulation of ECC mechanisms, and different propensity for development of Ca^2+^ instabilities.

**Figure 3.**
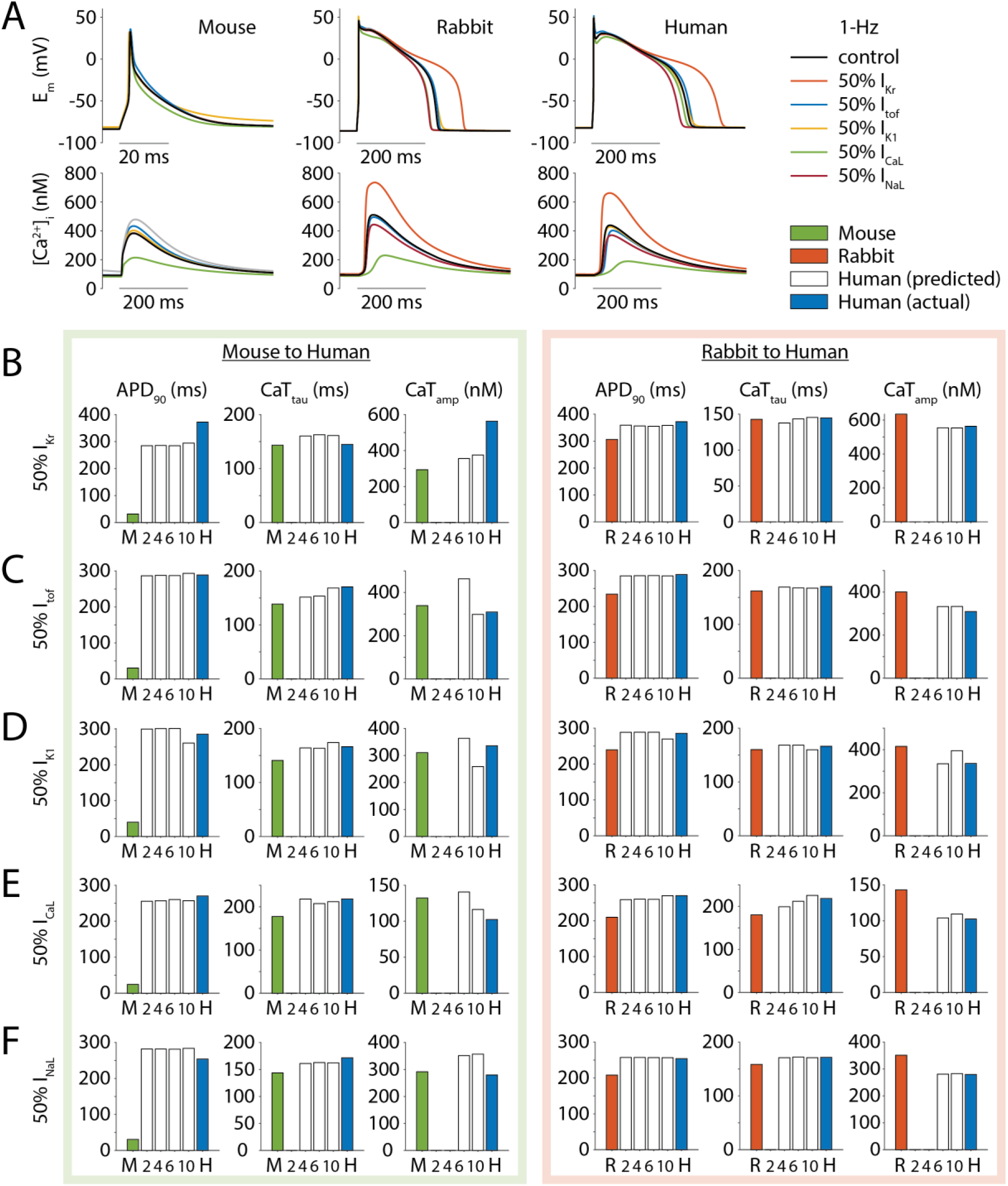
Validation of cross-species prediction against data simulating the effects of selective ion channel block. **A**) Simulated AP and CaT traces elicited when pacing the mouse, rabbit, and human models at 1-Hz in control condition or upon selective block (50%) of several ion currents. **B**-**F**) Bar graphs show AP and CaT features obtained in mouse (M, green), rabbit (R, orange), and human (H, blue) simulations, and predicted human data (white) obtained by applying the mouse-to-human (left panels) or rabbit-to-human (right panels) predictors built using a different number of features *n* (as described in **Fig. 2C**). Note that predictors built with two features can only predict APD_90_ and APD_50_ data (*n* = 2), and predictors built with four features can only predict APD_90_, APD_50_, CaT_tau_, and CaT_t50_ data (*n* = 4).

### Translation of measured drug-induced effects demonstrates robust predictions across species

As a next critical step, we sought to validate our translators against experimental data. To do so, we collected a set of published measurements describing the effects of ion channel blockers on APD. To account for the large degrees of experimental variation in these measures, we used the experimentally observed drug-induced relative changes to scale the APD values produced by our baseline models in control condition for both input and output species. We applied our previously described predictors (i.e., those built considering APD_90_ and APD_50_ data in control condition, *n* = 2) to the scaled input values, and then compared the results with the scaled output values for validation. Predictions of human response to the specific I_NaL_-blocker GS-967 from mouse^37^ and rabbit data^38^ closely reproduced the effect seen in human experiments (**Fig. 4A**).^39^ The same was true for the translations in the opposite direction (from human to mouse or rabbit) and those between mouse and rabbit (**Fig. S5A**). Translation of the effect induced by I_Kr_-block with administration of E-4031 (**Fig. 4B**)^40–42^ and Sotalol (**Fig. 4C**)^43^ predicted APD changes within the range of experimental variability in human, as also observed when predicting rabbit responses from human data (**Fig. S5B-C**). Translating the effects of selective block of I_K1_^40,44^ and I_CaL_^43^ also yielded good predictions (**Fig. S6**). These results show that translators built using minimal datasets (APD_90_ and APD_50_ data only) can robustly predict human responses from animal data.

**Figure 4.**
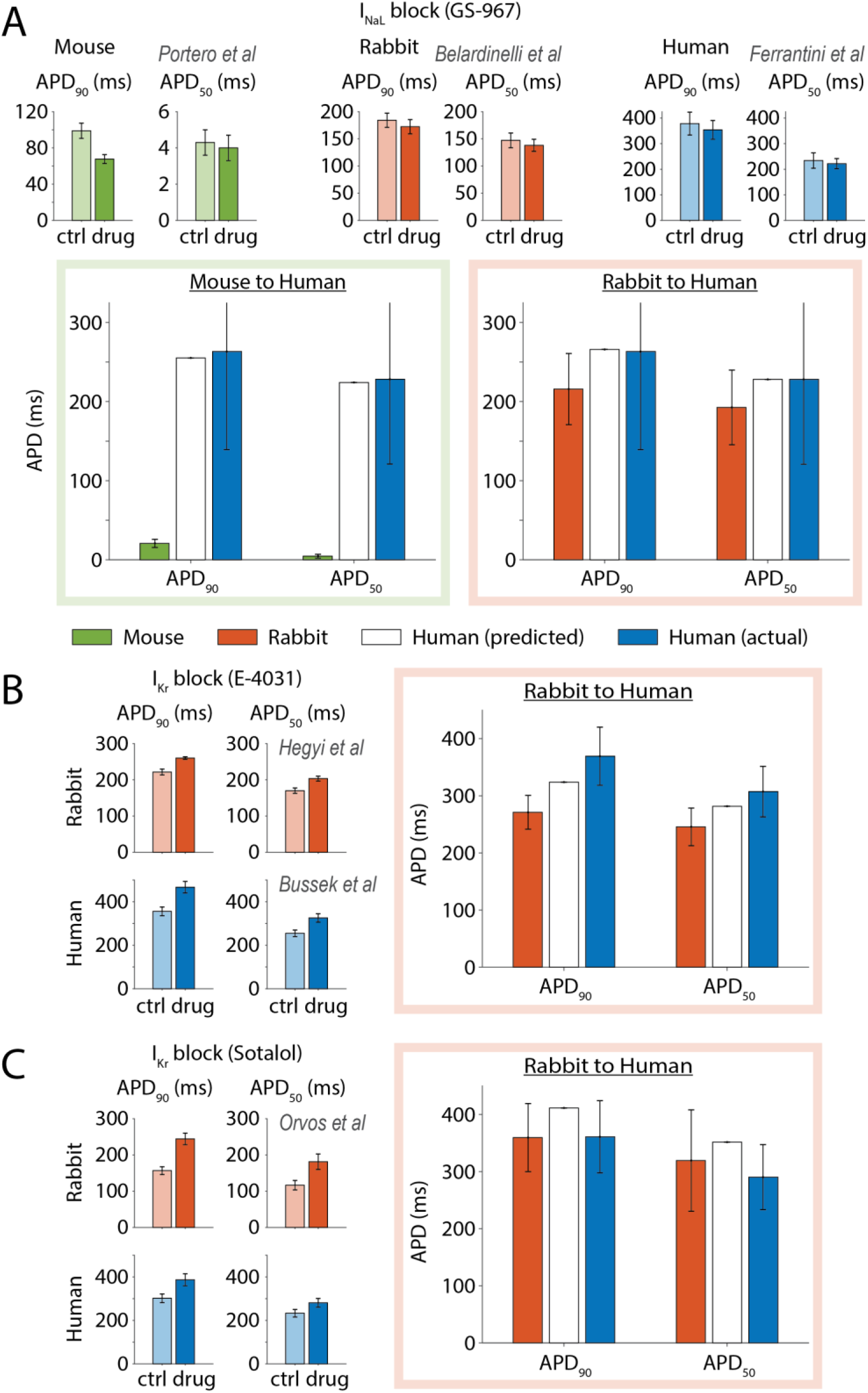
Experimental validation of cross-species prediction against experimental ion channel block data. **A**) The top panels illustrate experimental observations of the effect of the I_NaL_-blocker GS-967 on AP waveform in mouse (0.3 μM),^37^ rabbit (1 μM),^38^ and human (0.5 μM)^39^ ventricular myocytes. The bar graphs on the bottom display the validation of mouse-to-human and rabbit-to-human translations: mouse, rabbit, and human data are reported in green, orange and blue, respectively, while predictions of human response are reported in white. Experimentally measured effects of the **B**) I_Kr_-blocker E-4031 (1 μM) on AP waveform in rabbit^40^ and human ventricular myocytes (experimental data from ^41^, estimated as in ^42^) and **C**) I_Kr_-blocker Sotalol on AP waveform in rabbit (52 μM) and human (30 μM) ventricular myocytes^43^ at left are compared with the results of rabbit-to-human translation at right: rabbit and human data are reported in orange and blue, respectively, while predictions of human response are reported in white. All predictions shown here were obtained using the mouse-to-human or rabbit-to-human translators built using only APD_90_ and APD_50_ data (*n* = 2 in **Fig. 2C**). Error bars for experiments represent standard error of the mean, while error bars in the validation panels represent standard deviation.

### Analyzing translation performance allows identification of the most informative subset of input features

We demonstrated that translators’ performance depends on the size of the group of features included in its construction. In general, increasing the number of input features improves the performance because of the larger amount of information available in the definition of the regression model. However, AP and CaT features are not independent variables, but are potentially correlated with each other. We assessed the changes in the overall performance of our translators with a varying number of input features to find correlation among features and/or redundancy in the dataset, and identify the least and most informative features for the prediction of a certain group of AP and CaT properties in another species or experimental condition.

Specifically, we applied a recursive feature elimination routine to the prediction of human APD_50_, APD_90_, CaT_tau_, and CaTamp from mouse or rabbit data (**Fig. 5**). Starting from our complete set of ten features, this routine progressively eliminates the least informative input variable. Mouse-to-human prediction shows a first drop in overall performance starting at the fourth iteration, and a second one after reducing the input dataset to three features. Performance of rabbit-to-human prediction is mildly altered until reducing the input dataset to three features. Interestingly, both analyses lead to the identification of the same optimal four-feature group, including three of the four output features of interest (APD_90_, CaT_tau_, and CaTamp) and the minimum diastolic potential (MDP) that replaces the output feature APD_50_, discarded at earlier iterations in both cases. This suggests that MDP data would be more informative than APD_50_ data for the quantitative cross-species prediction of human APD_50_, APD_90_, CaT_tau_, and CaTamp (when APD_90_, CaT_tau_, and CaTamp animal data are available). The approach presented here might be used to guide the design of experiments on animal models, i.e., by identifying variables that are critical for accurate translation.

**Figure 5.**
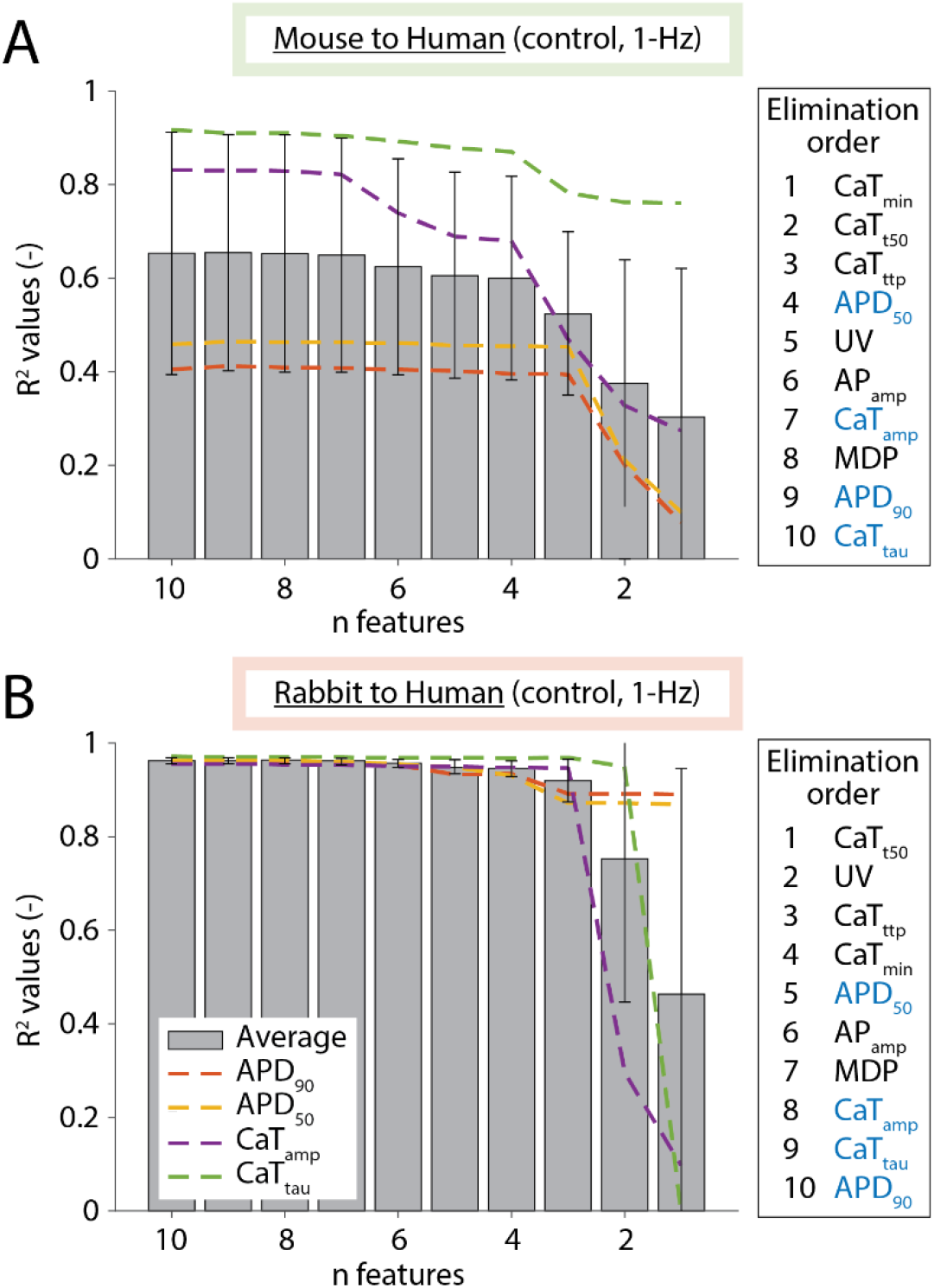
Assessment of input features’ informative score. Average R^2^ values characterizing the overall performance of mouse-to-human (**A**) and rabbit-to-human translators (**B**) built using a fixed number of output features (APD_90_, APD_50_, CaT_amp_, and CaT_tau_) and a variable number of input features. Starting from a set of ten input features, at each iteration an automatic recursive features elimination routine excluded the least informative input feature. Elimination order is reported on the right, where the features highlighted in blue are those produced in output for human. Error bars indicate standard deviation.

### Translation of measured responses to sympathetic stimulation demonstrates robust predictions across species and pacing rates

Our translators proved effective in predicting human response to ion channel block at a given pacing cycle length from animal data. A more challenging application of cross-species translation is in the context of sympathetic stimulation. First, β-AR stimulation functionally affects many subcellular targets at the same time (via activation of the protein kinase A, PKA, as shown in **Fig. S1**), thus affecting myocyte function in a multifaceted manner rather than with a single-targeted ion channel blocker. Second, sympathetic stimulation *in vivo* induces concomitant changes in sinoatrial node activity that result in increasing stimulation rates for the ventricular myocytes.

To decompose this complex problem, we first tested our predictions across different pacing rates against experimental data describing the frequency-dependence of rabbit APD_50_ and APD_90_ upon selective block of I_NaL_, I_to_, I_Kr_, I_Ks_, I_K1_ (**Fig. 6**), and the small-conductance Ca^2+^-activated K^+^ current (I_K,Ca_, **Fig. S7**).^40^ We generated simulated data with our population of rabbit models paced at different frequencies in control condition, and then built cross-frequency translators to predict responses at 0.5, 2, and 3 Hz from observations at 1 Hz. Cross-frequency translation produced reliable predictions when applied to drugs that significantly impact APD, causing both shortening (**Fig. 6A**) or different degrees of prolongation (**Fig. 6B-E**). Since apamin administration does not significantly alter (undiseased) rabbit ventricular AP, cross-frequency translation of I_K,Ca_ block essentially operates on control values, showing optimal results (**Fig. S7**).

**Figure 6.**
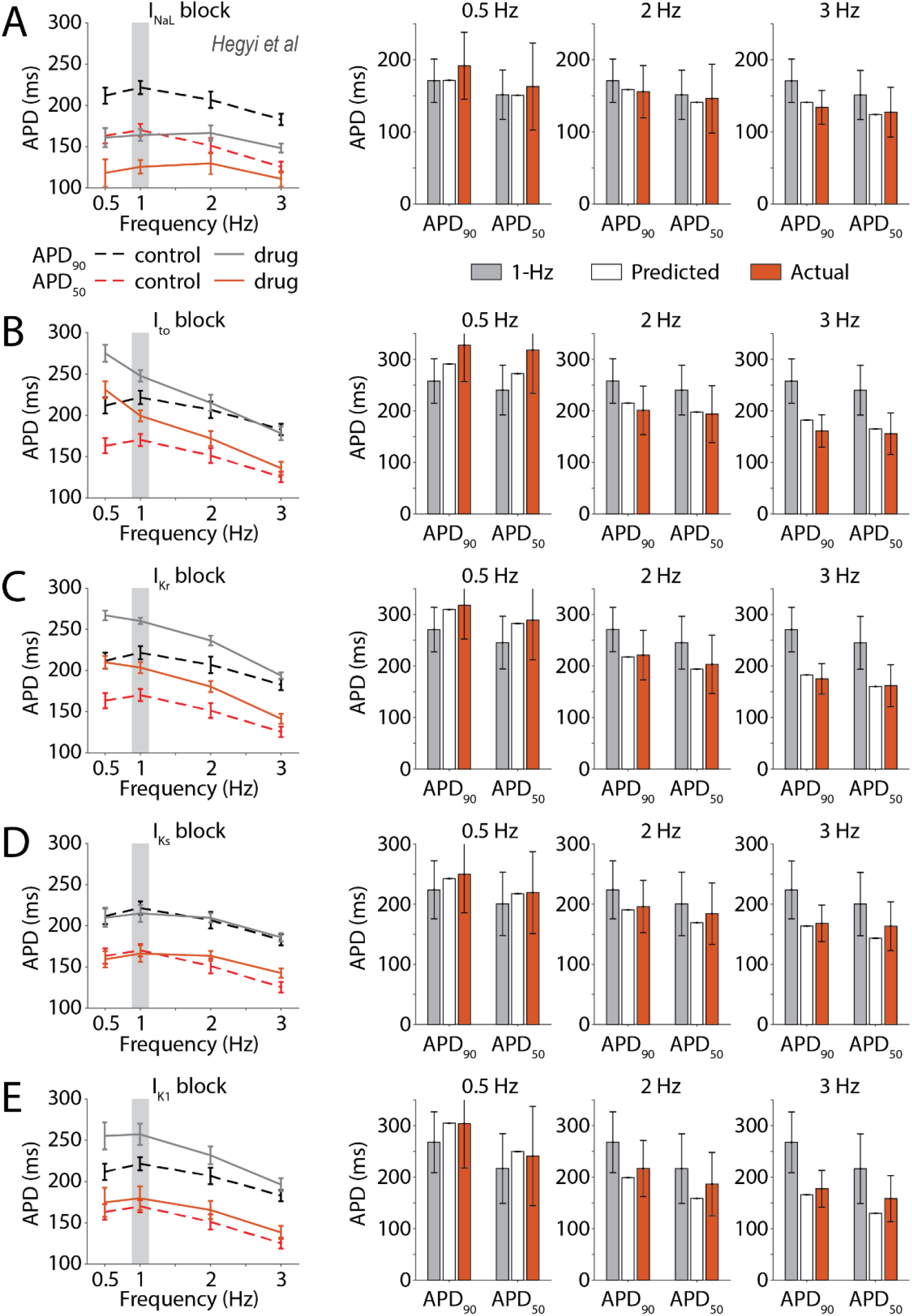
Experimental validation of cross-frequency prediction against data describing effect of ion channel block in rabbit myocytes. At left are changes in APD_90_ and APD_50_ induced by block of I_NaL_ (1 μM GS-967, panel **A**), I_to_ (5 mM 4-AP, **B**), I_Kr_ (1 μM E-4031, **C**), I_Ks_ (1 μM HMR-1556, **D**), and I_K1_ (200 nM PA-6, **E**) in rabbit myocytes paced at different frequencies.^40^ At right is the validation of cross-frequency translation of APD values after drug administration: 1-Hz data are reported in grey, while 0.5, 2, and 3-Hz data are in orange, and their predictions from 1-Hz data are in white. The predictors used here were built with APD_90_ and APD_50_ data (*n* = 2) obtained stimulating the population of rabbit models in control condition at the different pacing rates. Error bars in the left panels represent standard error, while error bars in the bar graphs represent standard deviation.

Next, we tested our translators against experimental data describing the effect of Isoproterenol (ISO) administration. We first applied our cross-frequency translators in rabbit to experimental data describing APD responses to ISO at various pacing frequencies (**Fig. 7A**).^40^ Prediction of APD effects at 0.5, 2, and 3-Hz pacing rate starting from 1-Hz data fell within the variability ranges observed in experiments in the three cases. We then tested cross-species prediction of ISO effect using our mouse^45^ and rabbit^40^ observations at 1-Hz pacing (**Fig. 7B**). We showed that prediction of rabbit response to ISO from mouse using only two APD inputs fell outside the ranges of experimental variability (**Fig. 7C-D**), but outcomes of mouse-to-rabbit translation improved when also considering CaT features as inputs. Translation from rabbit to mouse shows a similar trend (*n* = 2, **Fig. S8A-B**), with more reliable predictions of APD_50_ vs. APD_90_. These cross-species predictions are remarkable considering that experimental results showed opposite ISO-induced effects on APD in the two species.

**Figure 7.**
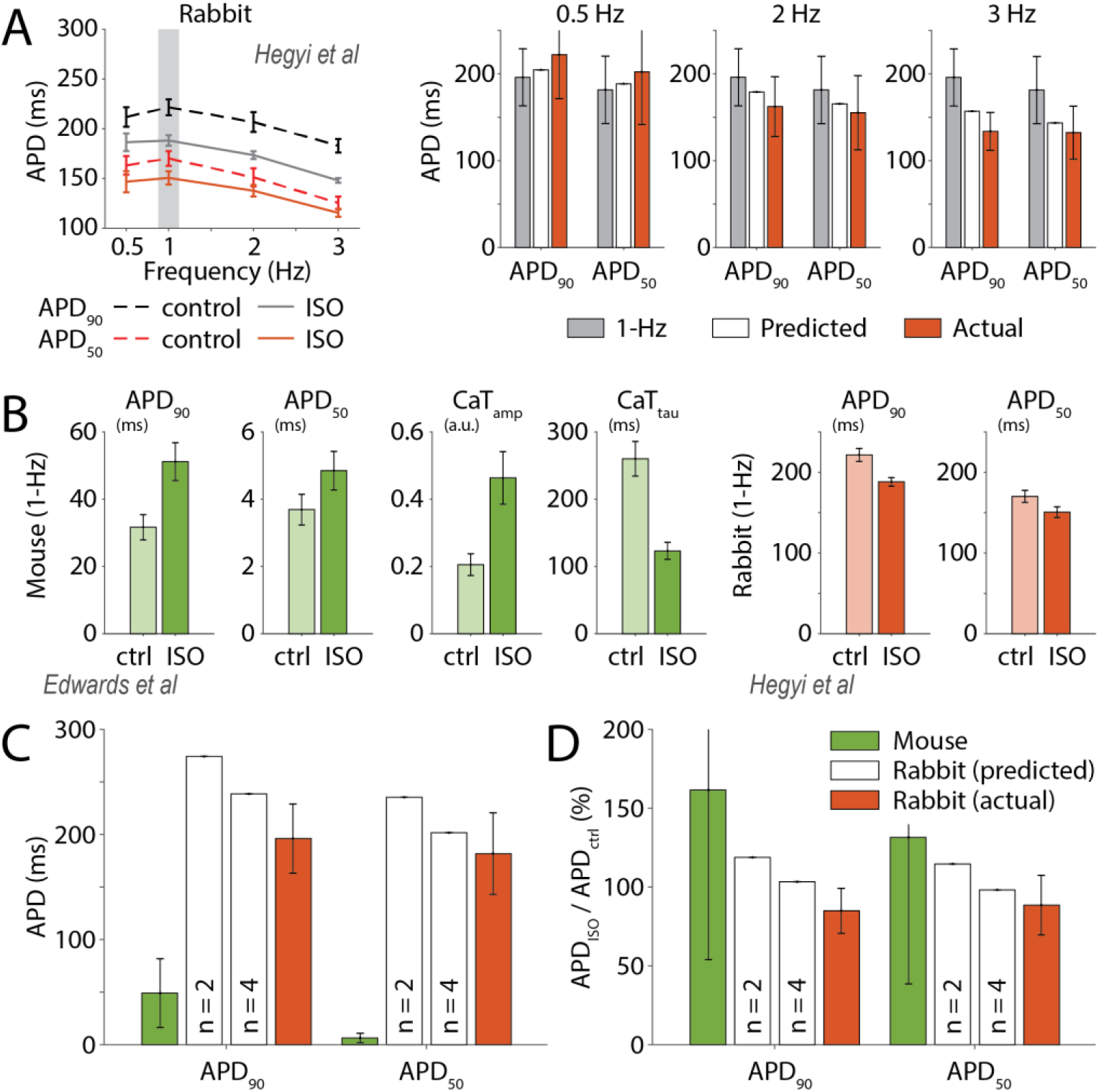
Experimental validation of prediction of Isoproterenol administration effects across pacing frequencies or species. **A**) At left are changes in APD_90_ and APD_50_ induced by ISO administration (30 nM) in rabbit myocytes paced at different frequencies.^40^ At right is the validation of cross-frequency translation of ISO effect: 1-Hz data are reported in grey, while 0.5, 2, and 3-Hz data are in orange, and their predictions from 1-Hz data are in white. The predictors used here were those described in **Fig. 6**. Error bars in the left panel represent standard error, while error bars in the bar graphs represent standard deviation. **B**) Experimental observation of the effect of ISO administration in mouse (100 nM)^45^ and rabbit (30 nM)^40^ ventricular myocytes. Error bars represent standard error. **C**) Mouse and rabbit data are reported in green and orange, respectively, while white bars represent prediction of rabbit response obtained with two mouse-to-rabbit translators built using different number of input features. The translator built with two features uses mouse APD_90_ and APD_50_ data only (*n* = 2), while the translator built with four features also uses mouse CaT_amp_ and CaT_tau_ data (*n* = 4). **D**) Actual and predicted ISO-induced APD changes estimated normalizing the values in panel **C** to the corresponding APD value assessed in the absence of ISO (i.e., control). Error bars in panel **C** and **D** represent standard deviation.

### Translators successfully predict cross-species effects of sympathetic stimulation in quasi-physiological conditions

In physiological conditions, sympathetic stimulation influences ventricular activity by combined PKA-dependent modulation of ion channels and transporter, and increased heart rate. To test cross-species translation in a condition more relevant for *in vivo* physiology, we used our recently published data describing the effect on APD and CaT duration (CaTD) recorded in innervated whole-heart mouse and rabbit preparations during sympathetic nerve stimulation (SNS) for 60 s (**Fig. 8A**).^22^ We built cross-species predictors that translate the relative effect induced by sympathetic activation using simulated data obtained by imposing the increases in stimulation rates seen in experiments in mouse and rabbit. Predicted relative effects induced by SNS on rabbit features nicely match experimental observations (**Fig. 8B**), as well as predictions of absolute APD and CaTD values after SNS (obtained scaling control rabbit data, **Fig. 8C**). Notably, the converse rabbit-to-mouse predictions are less accurate for CaTD (**Fig. S8C-D**). These results demonstrate the ability of our predictors to map complex responses to physiologically relevant stimuli across species.

**Figure 8.**
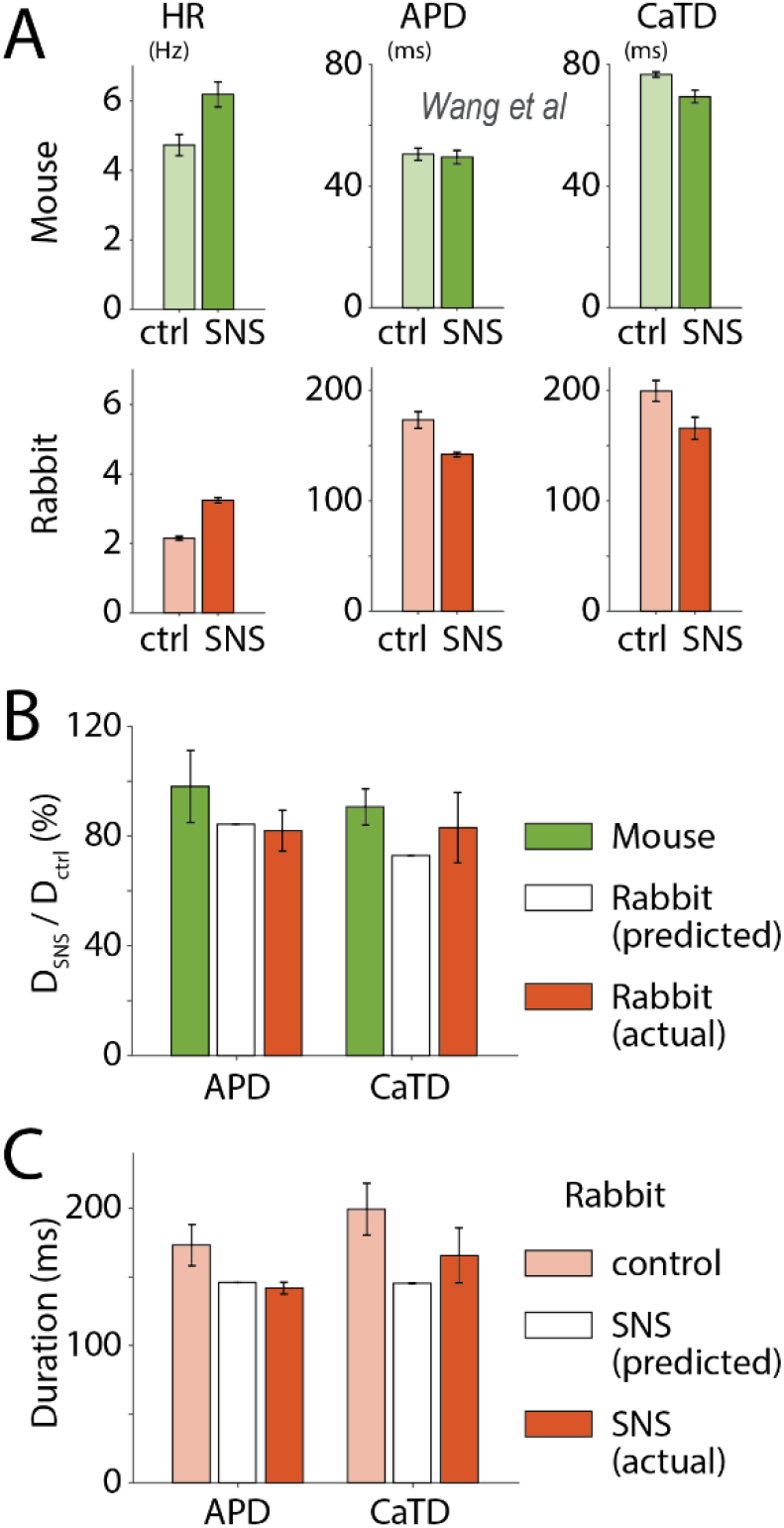
Experimental validation of cross-species prediction of sympathetic stimulation effect in quasi-physiological conditions. **A**) Experimental observation of the effect on heart rate (HR) and duration (D) of AP and CaT induced by SNS in mouse and rabbit innervated whole-heart preparations.^22^ Error bars represent standard error. **B**) Experimental validation of mouse-to-rabbit translation of SNS-induced relative APD and CaTD changes. Experimental mouse and rabbit data are reported in green and orange, respectively. Predicted rabbit response (in white) is obtained by applying a predictor built using relative changes in APD and CaTD estimated from simulations in mouse and rabbit populations that mimic the conditions observed in experiments. **C**) The relative SNS-induced effect in rabbit predicted form mouse data is used to estimate AP and CaT duration during SNS from control rabbit data. Error bars in panels **B** and **C** represent standard deviation.

## Discussion

We created a suite of regression-based operators to quantitatively translate electrophysiological responses across species,^29^ based on our updated models of mouse,^30,31^ rabbit,^32,33^ and human ventricular myocytes.^34,35^ We tested our translators against experimental data and demonstrated that these tools are well suited for predicting the human electrophysiological changes in response to a given stimulus (e.g., ion channel block and sympathetic stimulation) starting from the measured response in animal models. We also showed that these regression-based models can be used to inform the design of experiments by identifying the set of AP and CaT properties to be measured to maximize translatability to human physiology.

### Computational approaches to investigate implications of inter-species differences

Despite the limiting differences with respect to human physiology, animal models will remain an essential tool for investigating arrhythmogenic mechanisms and assessing safety and efficacy of new drugs. Thus, while pursuing a better characterization of these differences is undoubtedly important, a priority for the scientific community is to apply the existing knowledge to the development of reliable methods for translating findings in animals into human physiology. Computational modeling and simulations, now commonly used in the drug development pipelines,^46,47^ have proven useful for integrating species-specific experimental data into comprehensive mathematical models of mouse,^30,48^ rat,^49^ guinea pig,^50^ rabbit,^51^ dog,^52^ and human ventricular myocytes.^34,53,54^ For many years, implications of inter-species differences have been investigated simulating these models,^22,44,55^ or integrating simulations into the experimental activity, as in the case of the dynamic clamp technique.^21,56^

Recently, more complex methodologies have been specifically developed to facilitate translation across species or cell types. Tveito and colleagues proposed an approach based on the experimentally-driven automatic identification of drug effects on ion channels and transporters in animal models, and simulation of these alterations in a human model.^42,57,58^ While this demands performing mathematical analysis and running a new simulation after each experiment, the methodology proposed by Gong & Sobie allows direct transformation of measured data.^29^ This reproducibility advantage, due to the fact that the combined use of population of models and statistical regression is required only when constructing a new translator, makes the methodology we used well suited for a straightforward integration into experimental practice.

Populations of models have been used for over a decade in cardiac electrophysiology, and led to many novel insights into physiological and pathophysiological variabilities (including in arrhythmia mechanisms),^59^ and variable responses to drug administration.^60,61^ Indeed, analysis of populations of cardiac AP models have contributed to our understanding of the relative roles of the underlying properties (model parameters) in modulating a given phenotypic trait or biomarker (i.e., sensitivity analysis), or revealing association of certain parameter ranges or properties with specific outcomes.^36,62^ Here, we utilized these methods to quantify inter-species differences in the regulation of AP and CaT (**Fig. S2**), and to develop direct translators of electrophysiological response. A specific methodological strength of our translators is that they are based on a lineage of species-specific models,^30–35^ based on the Shannon *et al* and the Soltis & Saucerman ventricular rabbit ECC^51^ and signaling^63^ models, which all share the same assumptions and structure, allowing virtually any model parameter in one species to be mapped onto another species. Indeed, while Gong & Sobie showed that an improvement in the translation accuracy could be obtained by extending the “input” simulated datasets,^29^ we demonstrate here that our regression-based translators can produce reliable predictions even when developed and applied to reduced experimental datasets.

### Prediction of human or rabbit response from mouse data

Rabbit is considered a more reliable model than mouse because of the more human-like AP profile and APD regulation mechanisms.^10^ Nevertheless, mice are extensively used for studying cardiac electrophysiology, investigating arrhythmogenic mechanisms, and assessing drugs’ cardiotoxicity.^7–9^ Here, we showed that rabbit-to-human translation is generally highly accurate, and that our translators can produce a reliable prediction of the effects induced in human by ion channel block starting from rabbit experiments (**Fig. 4** & **Fig. S6**). We also showed that prediction of human response from mouse data is generally less accurate, especially for APD values. These results are not surprising considering the intrinsic differences in the mechanisms regulating the short and triangular AP in small rodents vs. larger mammals, as also highlighted by our sensitivity analysis (**Fig. 1D**). Moreover, different propensity for voltage or Ca^2+^ instabilities (as shown in **Fig. S4**) contributes to making the translation of mouse response even more challenging.^13^ While these observations support the general notion that experiments in rabbit are more informative than those in mouse, they also suggest that development of reliable translators of electrophysiological response is particularly important for mouse data. Notably, we showed that our translators of mouse data provide a reliable prediction of the effects on APD induced by block of I_tof_, I_K1_, I_CaL_, and I_NaL_ in human (**Fig. 3** & **Fig. 4A**), or induced by ISO administration in rabbit (**Fig. 7B-D**). The latter result is of particular interest because of the many factors influencing ventricular ECC during β-AR challenge, when even small differences in AP shape have been shown to contribute to a different APD response.^21^ Indeed, while β-AR stimulation directly induces cAMP-dependent activation of PKA (which in turn phosphorylates multiple protein targets within the cell, altering their function), at the same time it also increases CaMKII activity via enhanced cytosolic Ca^2+^ signal, leading to further functional modulation of the targets of CaMKII-dependent phosphorylation (often already affected by PKA, as shown in **Fig. S1**). Lastly, it is important to remark that the regression-based models used in these applications were built using “control” data (i.e., not obtained by simulating a specific drug action), and thus these tools could be applied to predict the effects of any possible perturbation (at a given pacing rate).

Characterizing drug response at different frequencies is fundamental for anti-arrhythmic agents aiming at exerting maximal effect at the fast pacing rates that characterize ventricular tachycardia and fibrillation. We showed that our translators can reliably predict electrophysiological response across stimulation frequencies for both ion channel block and ISO administration (**Fig. 6** & **Fig. 7A**) and could therefore be applied to assess the rate-dependent effect of new compounds limiting the number of wet experiments needed. The ability to make reliable predictions across frequencies is also important for successful mouse-to-rabbit translation of the effects of sympathetic stimulation in quasi-physiological conditions (**Fig. 8**). Indeed, this was a challenging task considering that, besides the direct effects of β-AR activation on ventricular cells described above, changes in the sinoatrial node (positive chronotropy) lead to different increases in stimulation rate in the different species.^22^ These observations suggest that adoption of our regression-based translators, built with our established models of ventricular electrophysiology and signaling pathways, could facilitate the development of anti-arrhythmic strategies based on the use of β-blockers.^23^

### Applications

We extended the hiPSC-CMs to adult myocyte translator originally developed by Gong & Sobie^29^ to build robust predictors that can reliably translate measured electrophysiological responses across different species or experimental conditions. In compliance with the 3R principles (replacement, reduction and refinement) in animal research,^64^ adoption of these tools in experimental practice could be beneficial in several applications of animal testing. For example, we envision a straightforward integration into high throughput systems used in the pharmaceutical industry to evaluate efficacy and safety of novel drugs. Furthermore, these regression-based models could be used to translate response from healthy to diseased conditions, as already proposed for heart failure,^29^ or across different stages of disease-induced remodeling, as seen in the progression of atrial fibrillation, or even to map age-related changes in electro(patho)physiology. Development of pharmacological approaches against atrial arrhythmias can also benefit from regional (e.g., atrial-to-ventricular) translators, to investigate potential adverse effects of compounds intended to selectively affect atrial function.^65^ Lastly, to overcome the negative implications of the vast underrepresentation of female sex in both basic research and clinical studies,^66^ translators of electrophysiological response across sexes could facilitate investigating differences in ECC regulation and arrhythmogenic mechanisms in male vs. female, and improve the assessment of drug cardiotoxicity in females.^67^

### Limitations and future directions

Although our baseline models have been developed and tuned in the years to reflect average properties observed in multiple studies from different groups, the experimental measures are quite variable. To account for this, when evaluating our translators against experimental data we translated the relative changes in the measured AP and CaT properties induced by a perturbation, rather than the absolute values. For any specific application, baseline models could be optimized to reproduce average features of the populations of cells used in experiments (sample-specific modeling) prior to constructing the translators.^68–70^

While the use one lineage of species-specific models helped improving translation accuracy, we acknowledge as an important implication the fact that these translators may reflect some specificities of our model lineage. Thus, future work should extend our approach to compare across other (and mixed) modeling lineages treated to a similar approach. Ideally, this major benchmarking process should be accompanied by the prospective design and collection of a broad experimental multi-species pharmacology dataset for assessing performance.

Lastly, our study focuses on ECC properties in the isolated ventricular myocyte. While AP and CaT dysregulations are important determinants, arrhythmia development and maintenance also critically depend on the characteristics of the cardiac tissue.^71^ Thus, important differences in organ dimensions and tissue-level properties (conduction velocity, effective refractory period, vulnerability window, and reentrant wave parameters) should also be taken into account when translating findings across species (or between healthy and diseased conditions) in the next generation of translators.

### Conclusions

Here, we used our updated multi-species framework for simulations of ventricular ECC to create a suite of robust cross-species translators of electrophysiological response. Our results suggest that integration of these regression-based models in both mechanistic studies and drug development pipelines will improve translation of findings in animal models by *i)* identification of more informative ECC features to be measured, and *ii)* direct prediction of the corresponding effect in human myocytes from experimental results. Extension of this approach to other cell types (such as hiPSC-CMs or atrial cells) or settings (health vs. disease, or male vs. female) could facilitate characterization of cardiac ECC physio-pathological mechanisms and development of safe and effective anti-arrhythmic strategies.

## Methods

### Updated multi-species framework for excitation-contraction coupling simulations in ventricular myocytes

All the simulations presented in this investigation were performed with our updated framework for simulating ECC in mouse, rabbit, and human ventricular myocytes (**Fig. S1**). Although their parameterizations differ to reflect species-specific observations, the three models share most components and have the same structure. In fact, membrane electrophysiology and intracellular Ca^2+^ and Na^+^ handling are modeled according to the geometry proposed for rabbit by Shannon *et al*,^51^ subsequently tuned to reproduce human-^34^ and mouse-specific properties.^30^ The common framework also includes detailed descriptions of CaMKII and β-AR/cAMP/PKA signaling pathways based on the work of Saucerman and colleagues.^63,72–74^ Changes to our previously published models are summarized below and in **Table S2**.

#### Baseline mouse model

We updated the CaMKII- and PKA-dependent signaling cascades in our model of the mouse ventricular myocyte.^30,31^ Namely, we included dynamic CaMKII-dependent regulation of I_NaL_;^75^ adjusted the dynamics and functional effect of PKA-dependent I_K,slow_ phosphorylation based on our previous observations;^22,76^ and added PKA-induced gain of function of the fast Na^+^ current (I_Na_),^48^ and loss of function of I_tof_ and I_K1_.^77,78^

#### Baseline rabbit model

Our model of the rabbit ventricular myocyte^32,33^ was modified to include CaMKII-dependent dynamic functional modulation of I_NaL_, as previously shown.^75^ As done in the initial development of our mouse model,^30^ the module describing β-AR activation and signaling cascade was transformed from a system combining algebraic and ordinary differential equations (ODEs) into a system with only ODEs. Modest adjustments to other model parameters were required to maintain the overall effect of sympathetic stimulation on AP and CaT (**Table S2**). PKA-dependent effect on I_Na_, I_tof_, and I_K1_ were included as well.^48,77,78^

#### Baseline human model

We modified our published model of the human ventricular myocyte^34,35^ by replacing the original I_Ks_ formulation with our updated version,^33^ and reducing its maximal conductance to maintain similar contribution to APD regulation. We also removed the cAMP-dependent (or Cystic Fibrosis Transmembrane Conductance Regulator) Cl^-^ current (I_CFTR_).^79^ As done for rabbit, the β-AR signaling cascade is now computed only with ODEs, and its functional effects are extended to I_Na_, I_tof_, and I_K1_.^48,77,78^ Lastly, we added the dynamic CaMKII-dependent regulation of I_NaL_.^75^

### Parameter sensitivity analysis

Parameter sensitivity analysis was performed with an established methodology based on the “populations-of-models” approach.^36^ We created species-specific populations of 1,000 models by randomly scaling the value of maximal conductance and transport rate of all ion currents and transporters (listed in **Fig. S2**) in each of our updated mouse, rabbit, and human models. For each model variant in the populations, the baseline value of each parameter was independently varied within a log-normal distribution (with σ = 0.1). As previously described,^61,80^ choice of variance and population size were such to ensure convergence of the sensitivity coefficients resulting from this analysis. AP and CaT properties (defined in **Table 1**) during 1-Hz pacing were assessed at steady-state in each model of the population. Variants showing inability of producing an AP, impaired repolarization, or alternans (2 models in the mouse population and 54 in the rabbit population) were excluded from further analysis. Multivariable regression analysis was performed to assess the influence of small variations in perturbed model parameters on AP and CaT features (**Fig. 1** & **Fig. S2**).^36^ The result of this process is the regression matrix B_SA_ that ensures X*B_SA_ ≈ Y, where X and Y are log-transformed parameter scaling factors and log-transformed AP and CaT properties, respectively. The matrix B_SA_ can be used to quantify the change in AP and CaT features upon perturbation in one or more model parameters.^36^

### Predictors of electrophysiological response

#### Generation and validation of the cross-species translators

To create and validate the translators, we built three additional populations of 1,500 models by randomly varying only the value of parameters common to the three species. Specifically, we did not alter *i)* I_Ks_ and the slow component of I_to_ (I_tos_), which are not expressed in mouse; *ii)* mouse-specific I_K,slow_ and I_ss_; *iii)* I_CFTR_, present only in the rabbit model (**Fig. S1** & **Table S1**). In this way, parameters in each model variant were varied according to the same matrix of scaling factors,^29^ as shown in **Fig. 1B**. Population size was here increased to create two groups to be separately used for creating and validating the translators. Specifically, we built each predictor using simulated data obtained in ~1,000-1,100 model variants (6 mouse and 72 rabbit models were excluded from the analysis because of AP abnormalities at 1-Hz pacing), and then tested their performance in the remaining and independent group of 400 *in silico* cells (next paragraph). For each model variant, AP and CaT features were assessed while pacing at different frequencies at steady-state in control condition, or after 60-s of 100 nM ISO administration. Following the methodology recently proposed by Song & Sobie,^29^ we created a group of predictors of electrophysiological response by applying multivariable linear regression on simulated log-transformed AP and CaT features. This process returns a regression matrix B_cross_ that ensures that Y_input_*B_cross_ ≈ Youtput, where Y_input_ and Youtput are log-transformed AP and CaT features obtained simulating different species (or experimental conditions). The matrix B_cross_ can be used to predict the value of AP and CaT features in the “output” species (or experimental condition) given the value of AP and CaT features in the “input” species (or experimental condition, **Fig. 1E**). Specifically, each output feature can be calculated applying a function in which each input feature is multiplied by its corresponding regression coefficient.^29^ We repeated the same process between different species, and between different experimental conditions (i.e., different pacing rates), varying the number of features considered.

To validate the translator, for each AP or CaT feature of interest, we compared the predicted values (obtained by applying B_cross_ to the data obtained simulating the input species/condition) and the actual values (obtained simulating the output species/condition), and calculated the coefficient of determination (R^2^). As overall performance index, we averaged the R^2^ values obtained for each feature.

We built a broad set of predictors by varying the composition of the groups of AP and CaT features considered in input and output. The rationale for reducing the number of features is that often only a subset of parameters is available from experiments, and we are interested in assessing the applicability of our translators to real data. We assessed the overall performance of translators using all ten AP and CaT features (*n* = 10) for both input and output (**Table 1**), and built on three subsets of features. As shown in the schematic in **Fig. 2C**, the minimal subset contained APD measurements only (APD_90_ and APD_50_, *n* = 2). In addition, intermediate subsets included measurements of CaT duration (CaT_t50_ and CaT_tau_, *n* = 4), and cytosolic Ca^2+^ concentration (CaT_min_ and CaT_amp_, *n* = 6).

Consequences of 50% block of I_tof_, I_Kr_, I_K1_, I_CaL_, I_NaL_, and NCX current on AP and CaT features were assessed at steady-state in our baseline mouse, rabbit and human models during 1-Hz pacing. The values of AP and CaT features assessed after ion channel block in mouse or rabbit were used as input for the prediction of AP and CaT features after ion channel block in human. We then compared predicted human values to actual values from human simulations for validation. This process was repeated using translators built with a variable number of features, as described above.

#### Experimental validation of the cross-species translators

An important validation aspect is to ensure that our translators can map experimental data measured in one species into data collected in another species. We mined the literature and used own experimental data to evaluate the ability of the translators to predict the effects of common perturbations on cardiac ECC, namely *i)* selective ion channel blockade, *ii)* changes in pacing frequency, and *iii)* effects of β-AR activation (ISO), which involve *i)* and *ii)*.

The relative perturbation-induced changes estimated from the experiments were used to scale the AP and CaT features predicted simulating the same species/condition with the corresponding baseline models in control condition (e.g., in the absence of ion channel block or ISO) for both input and output species/conditions. The “scaled” input features were translated to the output species/condition using the predictors (e.g., mouse-to-human at 1 Hz, or rabbit-1-to-3-Hz) built using control data obtained simulating the input species/condition (e.g., mouse at 1 Hz, or rabbit at 1 Hz) and the output species/condition (e.g., human at 1 Hz, or rabbit at 3 Hz). The resulting predicted values were then compared to the “scaled” output features for validation. Ranges corresponding to variability in scaled results (standard deviation) were estimated from experimental studies as well.

To perform experimental validation using our data describing the effects of SNS on APD and CaTD in innervated whole-heart mouse and rabbit preparations,^22^ we created cross-species translators of the relative effect induced by sympathetic activation in mouse and rabbit. To mimic the chronotropic effects seen in experiments, we simulated acute 100 nM ISO administration for 60 s, while increasing the pacing rate from 4.8 to 6.2 Hz and from 2 to 3.5 Hz for mouse and rabbit, respectively. For each element in the mouse and rabbit populations, we determined the relative changes in the duration of AP and CaT, then performed regression analysis between the two population matrices of relative changes. The resulting matrix B_cross_ can predict the relative SNS-induced change in the output species given the relative change in the input species. APD and CaTD values experimentally measured in the output species in control can then be scaled by the predicted amount and compared to the values measured during SNS for validation.

#### Performance analysis with recursive feature elimination

A recursive feature elimination routine was implemented to identify the more informative input features (among all ten of them) for the prediction of a fixed group of output features (APD_90_, APD_50_, CaT_amp_, and CaT_tau_). At each iteration, given a number *n* of available input features, this routine determines the overall performance (average R^2^) of the *n* predictors built ignoring one of the *n* input features at the time, and then discards the feature which elimination allowed for the best average R^2^. The routine stops when only one input feature remains.

### Code availability

All the codes used to perform simulations and data analysis were generated in MATLAB (The MathWorks, Natick, MA, USA), version R2018a. Population-level simulations were performed with a computing cluster with Intel(R) Xeon(R) CPU E5-2690 v4 @ 2.60GHz 28 CPUs (56 threads) + 132GB, and a standard laptop was used for data analysis. All our source codes (and related documentation) and all simulated data used in this study are available for download at http://elegrandi.wixsite.com/grandilab/downloads and https://github.com/drgrandilab.

## General

We thank Drs. Jakub Tomek and Jonathan Moreno for helpful discussion. We would also like to offer special thanks to Dr. Jorge Negroni, who, although no longer with us, continues to inspire our work.

## Funding

This work was supported by NIH/NHLBI Grants R00HL138160 (S.M.), R01HL131517 (E.G.), P01HL141084 (E.G. and D.M.B.), R01HL111600 (C.M.R.); NIH Stimulating Peripheral Activity to Relieve Conditions Grant 1OT2OD026580-01 (E.G.); American Heart Association Scientist Development Award 15SDG24910015 (E.G.), and Postdoctoral Fellowship 20POST35120462 (H.N.); UC Davis School of Medicine Dean’s Fellow Award (E.G.); Health and Environmental Sciences Institute Grant U01FD006676-01 (A.G.E.).

## Author contributions

S.M. and E.G. designed the research. S.M., C.L., and H.N. updated the baseline myocyte models and produced the simulated data. B.H., L. W., C.M.R., and D.M.B. provided experimental data. S.M., H.N., A.F.I., A.G.E., and E.G. analyzed the data. S.M. and E.G. wrote the manuscript, then revised by all other co-authors.

## Competing interests

The authors declare no competing interests.

## Data and materials availability

All data needed to evaluate the conclusions in the paper are present in the main text and the Supplementary Materials. Data and source codes are freely available for download at https://github.com/drgrandilab and http://elegrandi.wixsite.com/grandilab/downloads.

## Notes

### Competing Interest Statement

The authors have declared no competing interest.

