## Supplementary Materials for "Quantitative cross-species translators of cardiac myocyte electrophysiology: model training, experimental validation, and applications"

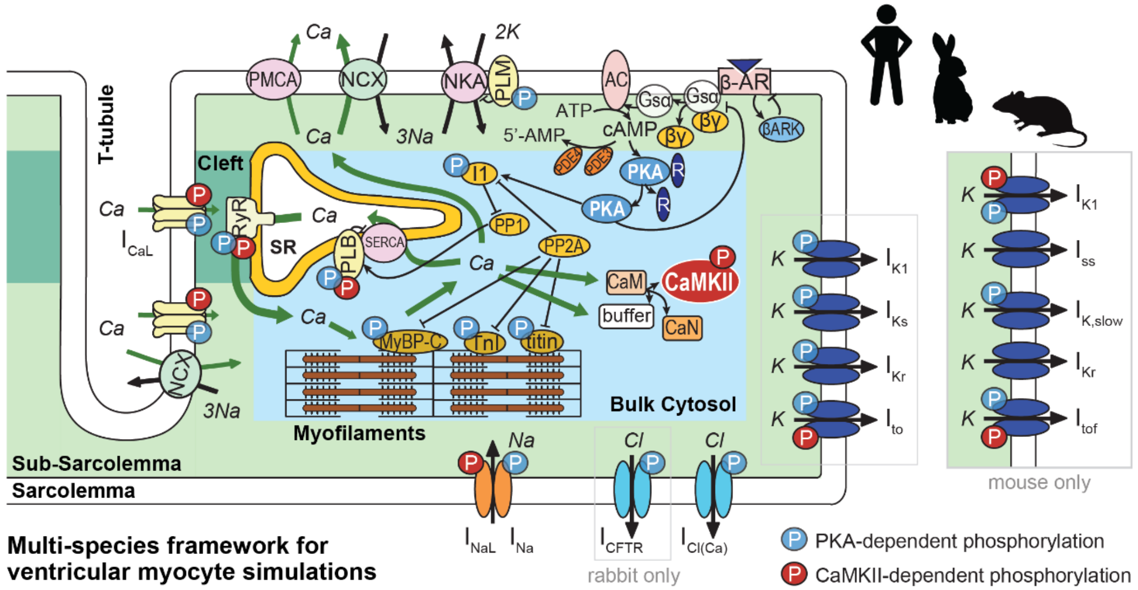


**Figure S1. Schematic of the computational modeling framework for mouse, rabbit, and human ventricular myocyte simulations.** Our updated multi-species modeling framework integrates detailed description of membrane electrophysiology, intracellular $Ca^{2+}$ and $Na^{+}$ handling, CaMKII and β-AR/cAMP/PKA signaling cascades, and myofilament contractility. The inset on the right highlights the different expression of $K^{+}$ channels in mouse ventricular myocytes.

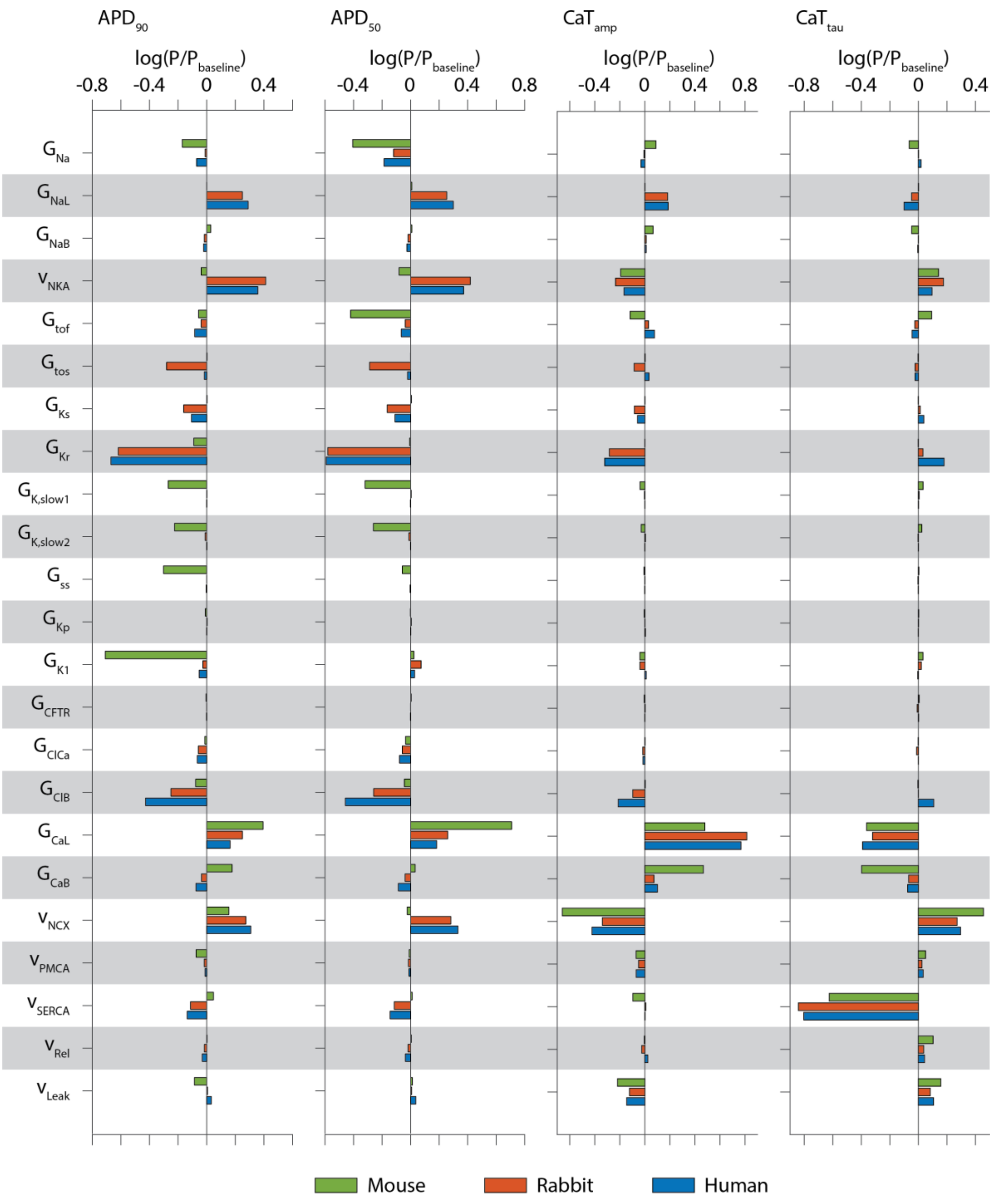


**Figure S2. Inter-species differences in the sensitivity of action potential and $Ca^{2+}$ transient features to changes in model parameters.** Regression coefficients quantifying the sensitivity of $APD_{90}$, $APD_{50}$, $CaT_{amp}$, and $CaT_{tau}$ (1-Hz pacing, control) to changes in ion channels' maximal conductance and transporters' maximal rate in the three species. Perturbed parameters are defined in **Table S1**.

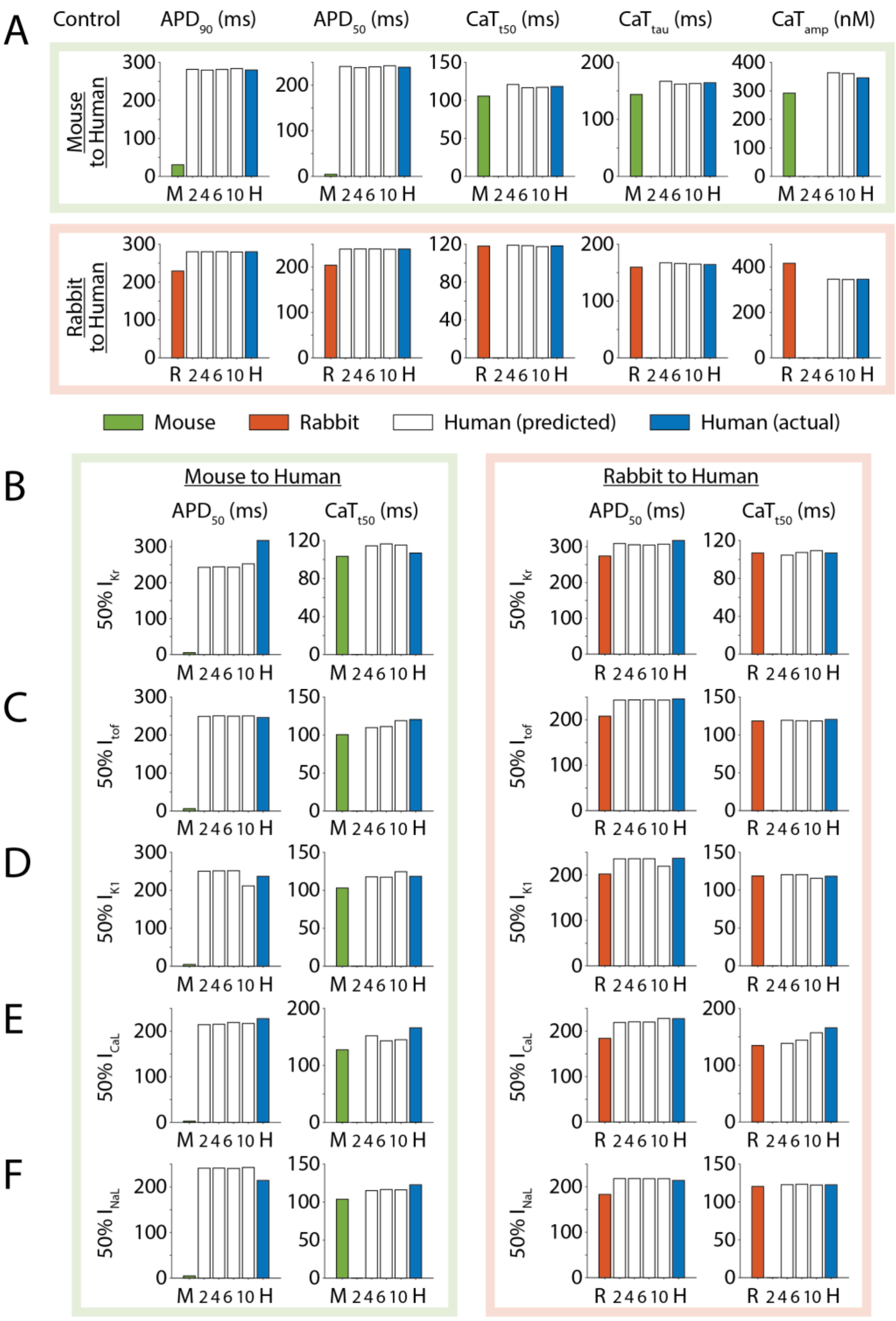


**Figure S3. Validation of cross-species prediction against simulated data describing control condition or effect of ion channel block. A**) AP and CaT features obtained pacing at 1 Hz the baseline mouse (M, green), rabbit (R, orange), and human (H, blue) models. **B**-**F**) $APD_{50}$ and $CaT_{t50}$ values obtained in mouse, rabbit, and human simulations while pacing at 1 Hz and blocking by 50% the indicated ion current. Predicted human data (white) were obtained applying the mouse-to-human or rabbit-to-human predictors built using a different number of features *n* (as described in **Fig. 2C**). Note that predictors built with two features can only predict

$APD_{90}$ and $APD_{50}$ data ($n$ = 2), and predictors built with four features can only predict $APD_{90}$, $APD_{50}$, $CaT_{tau}$, and $CaT_{t50}$ data ($n$ = 4).

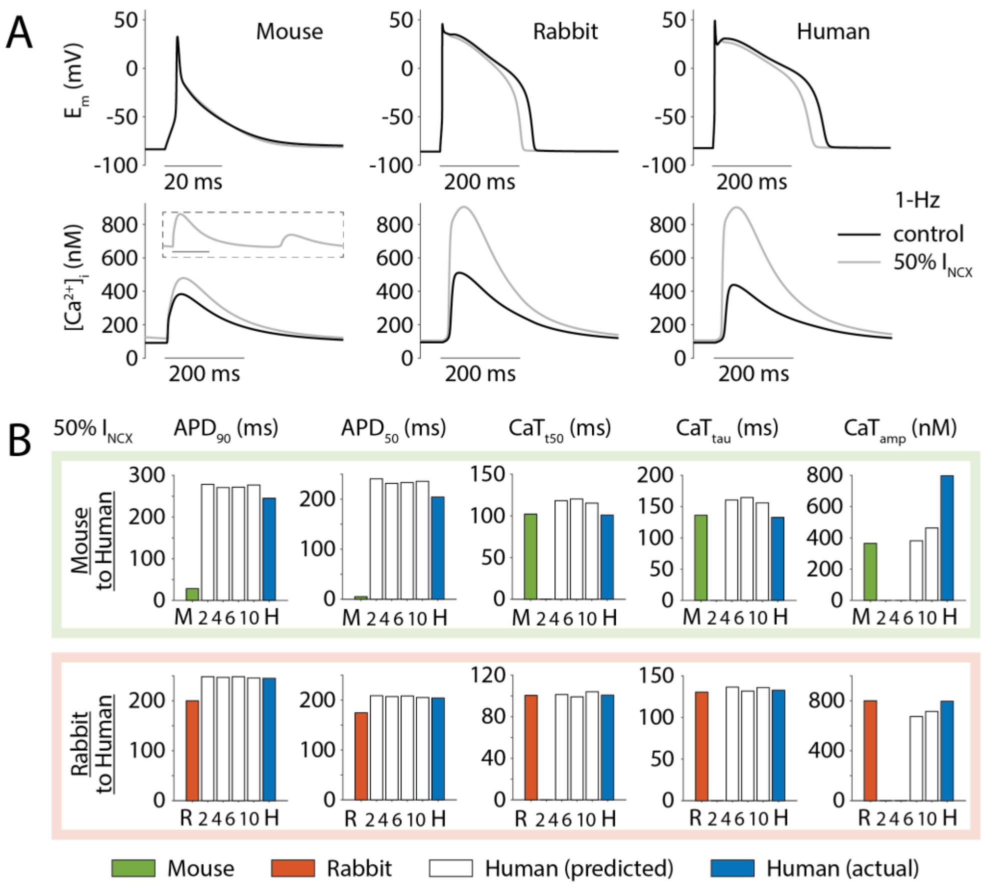


**Figure S4. Validation of cross-species prediction against simulated data describing the effect of selective block of $Na^+/Ca^{2+}$ exchanger. A**) AP and CaT traces elicited stimulating the baseline mouse, rabbit, and human model during 1-Hz pacing in control condition (black), or when selectively blocking NCX by 50% (grey). The inset shown in mouse CaT panel reports the trace elicited simulating NCX block over the whole stimulation period of 1 s (the vertical axis goes from 0 to 500 nM). **B**) AP and CaT features obtained in mouse (M, green), rabbit (R, orange), and human (H, blue) simulations, and predicted human data (white) obtained applying the mouse-to-human (top panels) or rabbit-to-human (bottom panels) predictors built using a different number of features $n$ (as described in **Fig. 2C**). Note that predictors built with two features can only predict $APD_{90}$ and $APD_{50}$ data ($n$ = 2), and predictors built with four features can only predict $APD_{90}$, $APD_{50}$, $CaT_{tau}$, and $CaT_{t50}$ data ($n$ = 4).

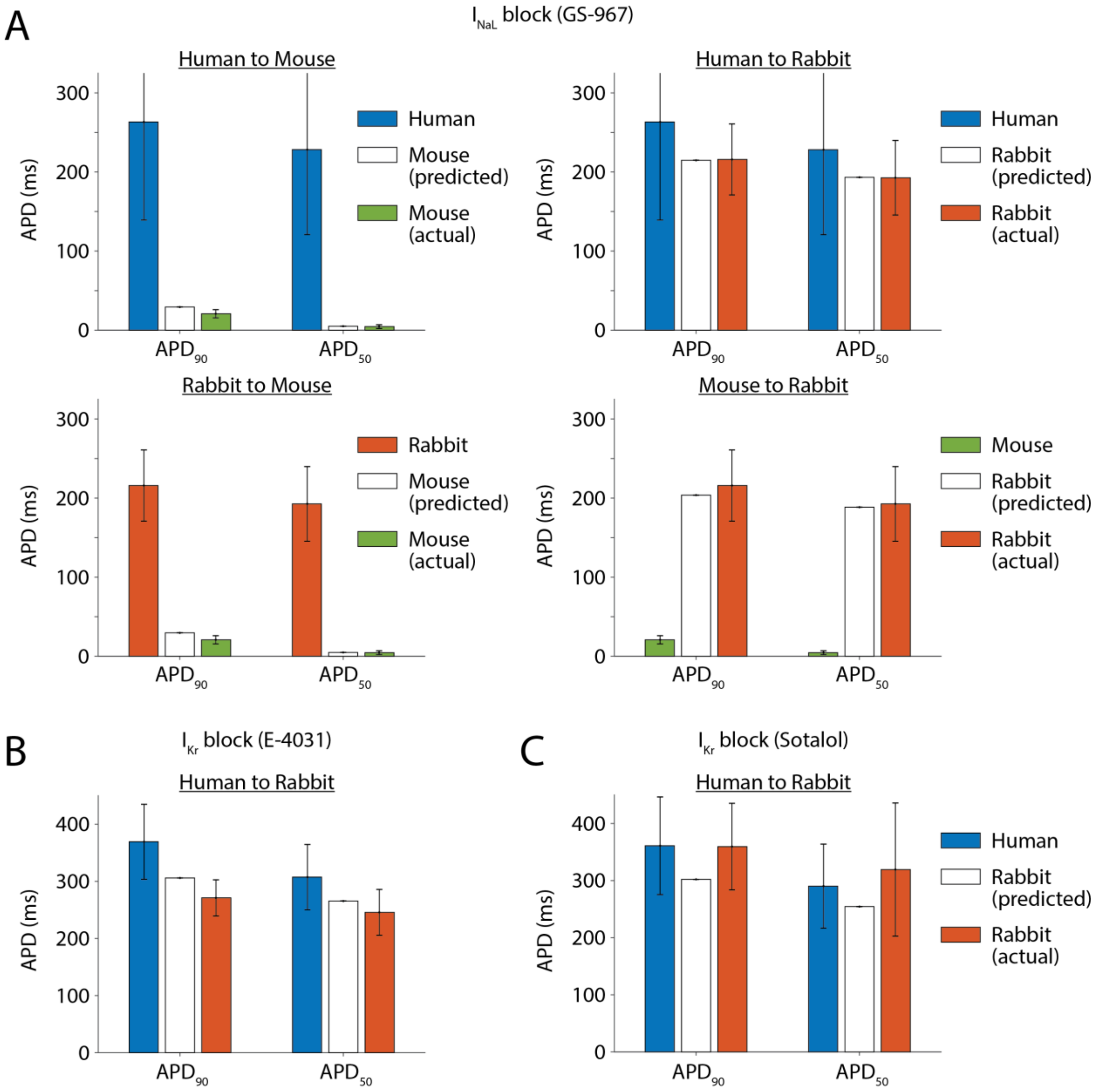


**Figure S5. Experimental validation of cross-species prediction against data describing effect of drug administration. A**) Cross-species prediction of the effect of the $I_{NaL}$-blocker GS-967. Top: validation of the human-to-mouse and human-to-rabbit translations. Bottom: validation of the mouse-to-rabbit and rabbit-to-mouse translations. **B**) Cross-species prediction of the effect of the $I_{Kr}$-blocker E-4031. **C**) Cross-species prediction of the effect of the $I_{Kr}$-blocker Sotalol. The graphs in panels **B** and **C** report the validation of the human-to-rabbit translations. All predictions shown here were obtained using translators built using only $APD_{90}$ and $APD_{50}$ data (*n* = 2). Error bars indicate standard deviation.

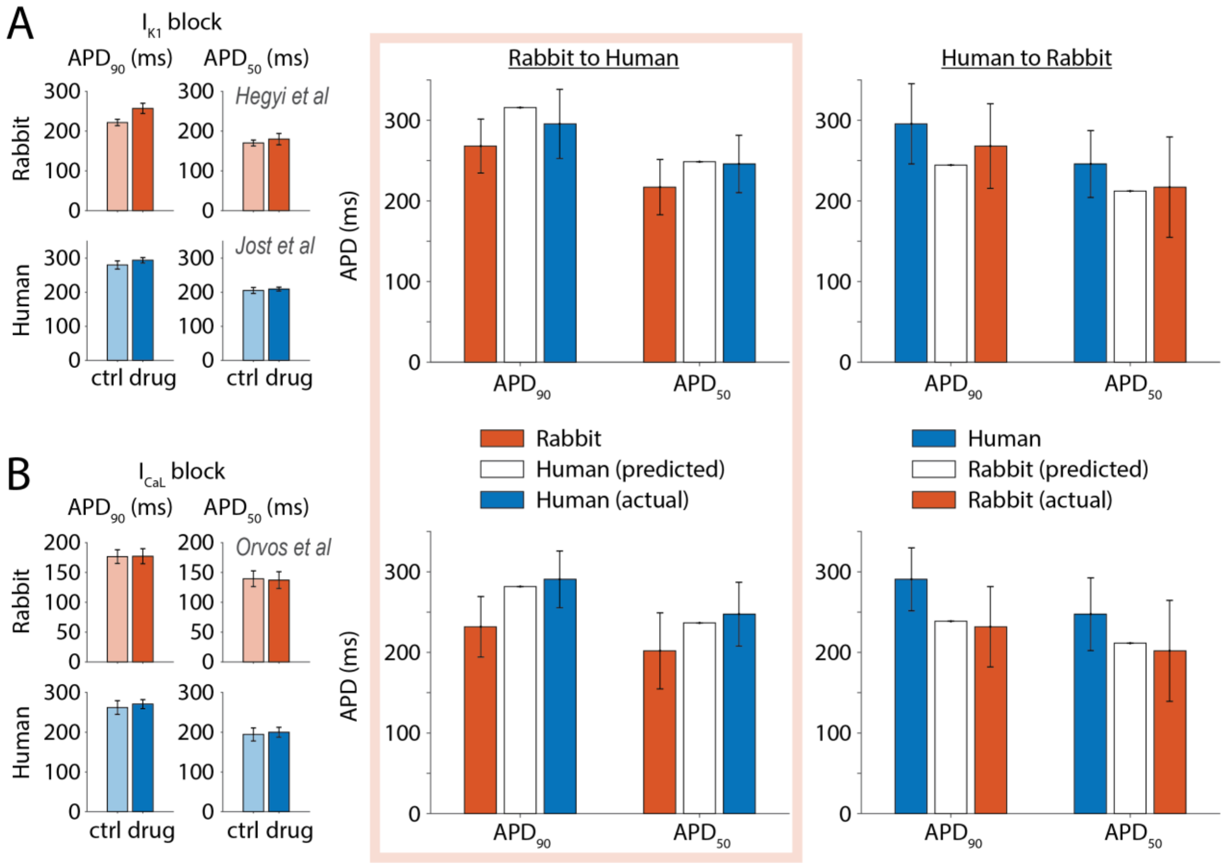


**Figure S6. Experimental validation of cross-species prediction against data describing effect of drug administration.** Left panels report experimental observations of **A**) the effect of selective $I_{K1}$ block on AP waveform in rabbit (200 nM PA-6)[40] and human (10 μM $BaCl_2$)[44] ventricular myocytes, and **B**) the effect of the $I_{CaL}$-blocker verapamil on AP waveform in rabbit (270 nM) and human (300 nM) ventricular myocytes.[43] At right is the validation of the rabbit-to-human and human-to-rabbit translations. All predictions shown here were obtained using the rabbit-to-human and human-to-rabbit predictors built using only $APD_{90}$ and $APD_{50}$ data ($n$ = 2). Error bars for experimental results (left panels) indicate standard error, while error bars in validation panels indicate standard deviation.

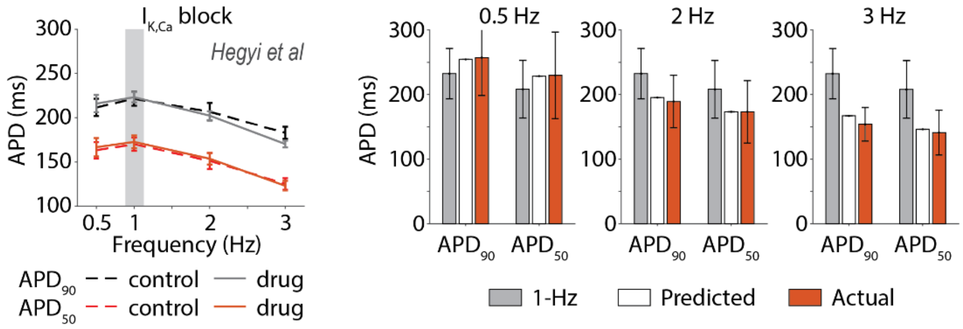


**Figure S7. Experimental validation of cross-frequency prediction against data describing effect of $I_{K,Ca}$ block in rabbit myocytes.** At left are changes in $APD_{90}$ and $APD_{50}$ induced by block of $I_{K,Ca}$ (100 nM apamin) in rabbit myocytes paced at different frequencies.[40] At right is the validation of cross-frequency translation of APD values after drug administration: 1-Hz data are in grey, while 0.5, 2, and 3-Hz data are in orange, and their predictions from 1-Hz data are in white. The predictors used here were those described in **Fig. 6**. Error bars in the left panel indicate standard error, while error bars in the bar graphs represent standard deviation.

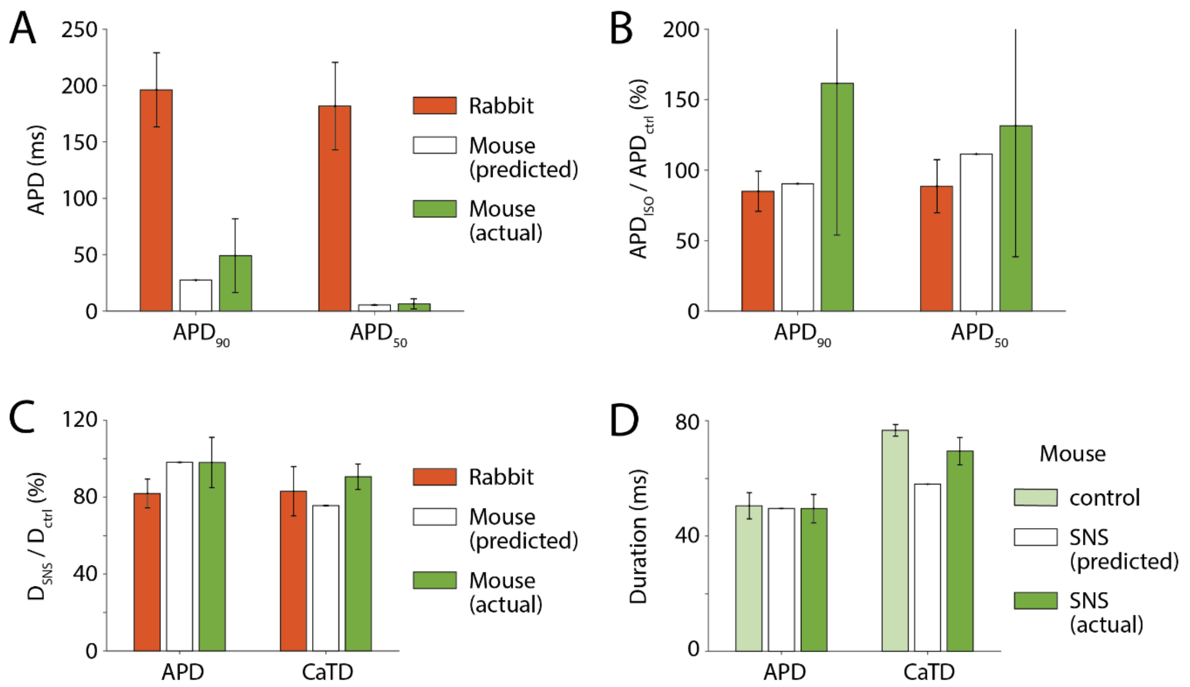


**Figure S8. Experimental validation of cross-species translation of the effect of sympathetic stimulation. A**) Validation of rabbit-to-mouse translation of ISO-induced effect during 1-Hz pacing, obtained applying a translator built using $APD_{90}$ and $APD_{50}$ data (*n* = 2). **B**) Actual and predicted ISO-induced APD changes estimated normalizing the values in panel **A** to the corresponding APD values assessed in the absence of ISO (i.e., control). Error bars indicate standard deviation. **C**) Experimental validation of rabbit-to-mouse translation of SNS-induced relative APD and CaTD changes in quasi-physiological conditions. Experimental mouse and rabbit data are reported in green and orange, respectively.[22] Predicted mouse response (in white) is obtained applying a predictor built using relative changes in APD and CaTD. **D**) The relative SNS-induced effect in mouse predicted form rabbit data in panel **C** is used to estimate APD and CaTD during SNS from control mouse data. Error bars indicate standard deviation.

**Table S1. Definition of model parameters varied to build the populations of models.** For each parameter, the letters at right indicate if description of the corresponding ion current or transporter is included into the models of mouse (M), rabbit (R), and human (H) ventricular myocytes.

| **Parameters** | | **Currents or Fluxes** | **Species** | | |
|---|---|---|---|---|---|
| $G_{Na}$ | Max conductance | Fast $Na^+$ current ($I_{Na}$) | M | R | H |
| $G_{NaL}$ | Max conductance | Late $Na^+$ current ($I_{NaL}$) | M | R | H |
| $G_{NaB}$ | Max conductance | Background $Na^+$ current ($I_{NaB}$) | M | R | H |
| $v_{NKA}$ | Max velocity | $Na^+/K^+$ ATPase current ($I_{NaK}$) | M | R | H |
| $G_{tof}$ | Max conductance | Transient outward $K^+$ current, fast component ($I_{tof}$) | M | R | H |
| $G_{tos}$ | Max conductance | Transient outward $K^+$ current, slow component ($I_{tos}$) | | R | H |
| $G_{Ks}$ | Max conductance | Slow delayed rectifying $K^+$ current ($I_{Ks}$) | | R | H |
| $G_{Kr}$ | Max conductance | Rapid delayed rectifying $K^+$ current ($I_{Kr}$) | M | R | H |
| $G_{K,slow1}$ | Max conductance | Ultra-rapidly activating and slowly inactivating $K^+$ current, 4-AP sensitive component ($I_{K,slow1}$) | M | | |
| $G_{K,slow2}$ | Max conductance | Ultra-rapidly activating and slowly inactivating $K^+$ current, TEA sensitive component ($I_{K,slow2}$) | M | | |
| $G_{ss}$ | Max conductance | Non inactivating steady-state $K^+$ current ($I_{ss}$) | M | | |
| $G_{Kp}$ | Max conductance | Plateau $K^+$ current ($I_{Kp}$) | M | R | H |
| $G_{Kp}$ | Max conductance | Inwardly rectifying $K^+$ current ($I_{K1}$) | M | R | H |
| $G_{CFTR}$ | Max conductance | cAMP-activated $Cl^-$ current ($I_{CFTR}$) | | R | |
| $G_{ClCa}$ | Max conductance | $Ca^{2+}$-activated $Cl^-$ current ($I_{ClCa}$) | M | R | H |
| $G_{ClB}$ | Max conductance | Background $Cl^-$ current ($I_{ClB}$) | M | R | H |
| $G_{CaL}$ | Max conductance | L-type $Ca^{2+}$ current ($I_{CaL}$) | M | R | H |
| $G_{CaB}$ | Max conductance | Background $Ca^{2+}$ current ($I_{CaB}$) | M | R | H |
| $v_{NCX}$ | Max velocity | $Na^+/Ca^{2+}$ exchanger current ($I_{NCX}$) | M | R | H |
| $v_{PMCA}$ | Max velocity | Plasma membrane $Ca^{2+}$ ATPase current ($I_{PMCA}$) | M | R | H |
| $v_{SERCA}$ | Max velocity | Sarcoplasmic reticulum $Ca^{2+}$ ATPase flux ($J_{up}$) | M | R | H |
| $v_{Rel}$ | Max velocity | Ryanodine receptor $Ca^{2+}$ flux ($J_{Rel}$) | M | R | H |
| $v_{Leak}$ | Max velocity | Ryanodine receptor $Ca^{2+}$ leak ($J_{Leak}$) | M | R | H |

**Table S2. List of changes in the updated baseline models.** Summary of the changes made to our previously published models of mouse, rabbit, and human ventricular myocytes.[30,33,35] Note that the β-AR modules in the rabbit and human models were transformed from a system combining algebraic differential equations and ODEs into a system with only ODEs.

| | Mouse | Rabbit | Human |
|---|---|---|---|
| ***Membrane electrophysiology*** | | | |
| $I_{CaL}$ | | 8% reduction in $G_{CaL}$ | |
| $I_{Kr}$ | | 40% increase in $G_{Kr}$ | 17% increase in $G_{Kr}$ |
| $I_{Ks}$ | | 40% increase in $G_{Ks}$ | Included new formulation[33] |
| $I_{ClB}$ | | 50% decrease in $G_{ClB}$ | |
| $I_{CFTR}$ | | | Removed[79] |
| ***β-AR signaling cascade & effects of PKA-dependent phosphorylation*** | | | |
| [PDE3] & [PDE4] | | 25% reduction | |
| SERCA | | Reduced (-36%) maximal PKA-dependent reduction of $k_{mf}$ | |
| $I_{CaL}$ | | Increased (+50%) maximal PKA-dependent enhancement of $G_{CaL}$ | |
| $I_{Na}$ | Included dynamic PKA-dependent regulation of $G_{Na}$: 8% increase with 100 nM ISO[48] | Included dynamic PKA-dependent regulation of $G_{Na}$: 25% increase with 100 nM ISO[48] | |
| $I_{to}$ | Included dynamic PKA-dependent regulation of $G_{tof}$: 13% increase with 100 nM ISO[77] | Included dynamic PKA-dependent regulation of $G_{tof}$: 40% reduction with 100 nM ISO[77] | |
| $I_{K1}$ | Included dynamic PKA-dependent regulation of $G_{K1}$: 15% reduction with 100 nM ISO[77,78] | Included dynamic PKA-dependent regulation of $G_{K1}$: 45% reduction with 100 nM ISO[77,78] | |
| $I_{K,slow}$ | Slower phosphorylation dynamics;[76] increased (5-fold) maximal PKA-dependent $G_{K,slow1}$ enhancement[22] | | |
| ***CaMKII signaling cascade & effects of CaMKII-dependent phosphorylation*** | | | |
| $I_{NaL}$ | Included dynamic CaMKII-dependent increase in $G_{NaL}$[75] | | |
